# HP1*β* and densely packed chromatin form separate microdomains in mouse ES cells, which are reconfigured upon exit from naïve pluripotency

**DOI:** 10.1101/2025.08.25.671447

**Authors:** Aleksandra Ochirova, Devina Shah, Ziwei Zhang, Aleks Ponjavic, Kazimir Uzwyshyn-Jones, David Lando, Meike Wiese, Xiaoyan Ma, Nicola Reynolds, Andria Koulle, Wayne Boucher, Martin O. Lenz, Brian D. Hendrich, David Klenerman, Ernest D. Laue

**Affiliations:** Department of Biochemistry, University of Cambridge, 80 Tennis Court Road, Cambridge CB2 1GA, United Kingdom; Department of Biochemistry and Molecular Cell Biology, Shanghai Key Laboratory for Tumor Microenvironment and Inflammation, Key Laboratory of Cell Differentiation and Apoptosis of Chinese Ministry of Education, Shanghai Jiao Tong University School of Medicine, Shanghai, China; Yusuf Hamied Department of Chemistry, University of Cambridge, Lensfield Road, Cambridge, CB2 1EW, United Kingdom; The Gurdon Institute, University of Cambridge, Tennis Court Road, Cambridge CB2 1QN, United Kingdom; Wellcome-MRC Cambridge Stem Cell Institute, Jeffrey Cheah Biomedical Centre, Puddicombe Way, Cambridge CB2 0AW, United Kingdom; Cambridge Advanced Imaging Centre, University of Cambridge, Downing Site, Cambridge CB2 3DY, United Kingdom; Department of Proteomics, The Novo Nordisk Foundation Center for Protein Research, Faculty of Health and Medical Sciences, University of Copenhagen, Blegdamsvej 3B, 2200 Copenhagen, Denmark; School of Physics and Astronomy, University of Leeds, Woodhouse Lane, Leeds, LS2 9JT, United Kingdom; School of Food Science and Nutrition, University of Leeds, Woodhouse Lane, Leeds, LS2 9JT, United Kingdom; UCL Cancer Institute, Paul O’Gorman Building, 72 Huntley St, London WC1E 6DD, United Kingdom; Department of Chromatin Regulation, Max Planck Institute of Immunobiology and Epigenetics, Stübeweg 51, D-79108 Freiburg, Germany; Department of Life Sciences, Imperial College London, London SW7 2AZ, UK

**Keywords:** Heterochromatin structure, heterochromatin protein 1, chromatin microdomains, liquid-liquid phase separation, ES cells, pluripotency exit, formative state, super-resolution imaging, single molecule tracking, histone H3K9 methylation

## Abstract

Heterochromatin proteins play a key role in establishing local chromatin structure to control the transcription of target genes. Here we uncover a surprising segregation between regions of high DNA- and high heterochromatin protein 1*β* (HP1*β*)-density in mouse ES cells. DNA-low/HP1*β*-high foci retain freely diffusing HP1*β*, and form via condensation through a multitude of weak interactions on top of HP1*β* that is bound stably to chromatin. DNA-high/HP1*β*-low foci exclude freely diffusing HP1*β* and display reduced chromatin mobility, suggesting a higher degree of chromatin self-interaction and a more repressive environment. Finally, the two types of environments are intertwined in DNA-high/HP1*β*-high foci, where HP1*β* maintains heterochromatin in a more compact yet dynamic chromatin state. During the exit from naïve pluripotency HP1*β* is lost from regions of high DNA density as cells transition through the formative state, which might facilitate the reconfiguration of genome structure accompanying a change in cell state that we observed previously. Subsequently, as cells enter primed pluripotency, canonical heterochromatin is established.

## Introduction

In a multicellular organism all cells contain the same genetic information but express a different subset of genes to give cells distinct identities and functions. The expression of cell type-specific genes at the transcriptional level is controlled by two competing processes: active transcription and the formation of heterochromatin. On the one hand, transcription factor binding to specific enhancers and promoters triggers entire transcriptional programs leading to changes in cell state, a paradigm beautifully illustrated by the conversion of fibroblasts to myoblasts through the expression of a single transcription factor MyoD (Davis *et al*, 1987). On the other hand, multiple different protein complexes function to establish heterochromatin structure through repressive post-translational modifications of histones (notably H3K9me2/3 and H3K27me3), leading to silenced states that can be maintained through cell division. A good example is heritable gene silencing in plants through H3K27 methylation and the Polycomb proteins (Yang *et al*, 2017), (Berry *et al*, 2017). In this study we have sought to further understand the structure and dynamics of heterochromatin, the principles of its formation and maintenance, and their influence on its repressive functions.

Whilst many of the mechanisms that define the features of heterochromatin have been revealed, a general model explaining their roles and interplay at different scales is still a matter of hot debate. The “classic” model of heterochromatin organisation depicts chromatin as a cross-linked globule, where some exogenous factor (such as HP1 – see below) acts as a staple linking distant regions together (Erdel & Rippe, 2018). This results in chromatin condensation and molecular crowding, which limits access to these regions by the bulky transcription machinery or other macromolecular complexes and thus provides the basis for transcriptional repression. Consistently, an inverse correlation exists between the size of the chromatin-modifying enzymes and the density of the chromatin compartments where they exert their function (Miron *et al*, 2020). On the other hand, it has been noted that the intrinsic self-interaction properties of chromatin are sufficient to drive chromatin precipitation under some conditions *in vitro*, especially when aided by DNA charge neutralisation by histone H1, and that these could well be relevant for heterochromatin condensation, its low mobility and accessibility in cells (Hansen *et al*, 2021).

A different view of heterochromatin as a more plastic and dynamic compartment than had previously been appreciated has emerged in recent years, initiated by the seminal finding that HP1 within heterochromatin of living cells exhibits surprisingly high mobility (Cheutin *et al*, 2003), (Festenstein *et al*, 2003). This view posits that heterochromatin might form via some form of liquid-liquid phase separation (LLPS), either inherent to the chromatin fiber itself (Gibson *et al*, 2019) or initiated by other proteins (Larson *et al*, 2017), (Strom *et al*, 2017), (Wang *et al*, 2019). This model implies that the accessibility or inaccessibility of heterochromatin might be conferred via the existence of a selective phase boundary – factors that interact with the constituents of the condensate are recruited, whilst the ones that do not are excluded (Banani *et al*, 2017), (Larson *et al*., 2017), (Strom *et al*., 2017), (Mittag & Pappu, 2022).

Heterochromatin protein 1 (HP1) is a hallmark protein of heterochromatin that is conserved from yeast to plants and mammals. In mammals, including mice and humans, there are three isoforms of HP1: HP1 *α*, HP1*β* and HP1 *γ*, which show different localisation patterns and perform partially overlapping, but distinct functions. All HP1 proteins consist of a chromodomain that interacts with the heterochromatic H3K9me2/3 histone modification (Bannister *et al*, 2001), a chromo shadow domain responsible for homo-dimerisation and the recruitment of a large variety of other protein complexes, and a flexible hinge connecting the two (Murzina *et al*, 1999), (Brasher *et al*, 2000). The HP1 dimer can bridge nucleosomes by binding to two histone H3 tails modified at Lysine-9 (Thiru *et al*, 2004), and consequently, HP1 is able to compact nucleosome arrays or even connect separate arrays *in vitro* (Hiragami-Hamada *et al*, 2016), (Fan *et al*, 2004), (Machida *et al*, 2018). Furthermore, HP1 has been shown to undergo LLPS *in vitro* either by itself or together with DNA, H3K9me3-modified chromatin and/or other heterochromatic proteins – with some evidence supporting and other studies challenging the physiological relevance of this process for heterochromatin formation in cells (Larson *et al*., 2017), (Strom *et al*., 2017), (Wang *et al*., 2019), (Erdel *et al*, 2020). In addition, HP1 recruits lysine methyltransferases that deposit the H3K9me2/3 modification on chromatin, thus propagating the heterochromatic state (Lechner *et al*, 2000), (Schultz *et al*, 2002), (Schotta *et al*, 2002), (Yamamoto & Sonoda, 2003), (Chin *et al*, 2007). Consistently, ectopic recruitment of HP1 to euchromatic loci results in gene repression, although the precise mechanism behind this is not known (Ayyanathan *et al*, 2003).

In this work, we studied heterochromatin and its relationship with the presence or absence of HP1*β* in mouse ESCs. We found that in these cells, heterochromatin is not always enriched in HP1*β*, and that often HP1*β* foci do not contain condensed chromatin. Furthermore, within foci, areas of high HP1*β* or of high DNA density tend to segregate away from each other. In DNA-poor/HP1*β*-rich foci ∼73% of the HP1*β* is bound to chromatin that is extensively H3K9me3-modified. These foci also retain freely diffusing HP1*β*, and we find evidence that they form via condensation through a multitude of weak interactions on top of HP1*β* that is bound stably to chromatin. On the other hand, DNA-high/HP1*β*-low areas exclude freely diffusing HP1*β* and display reduced chromatin mobility, suggesting a higher degree of chromatin cross-linking and a more repressive environment. Finally, in DNA-high/HP1*β*-high foci, the two types of environments are intertwined, leading to a more compact but dynamic chromatin state. Importantly, upon exit from naive pluripotency, the enrichment of HP1*β* within heterochromatin is first decreased, followed by the formation of more “canonical” heterochromatin with a much higher degree of overlap between HP1*β*-high and DNA-high regions. Our results suggest that HP1*β* functions to keep chromatin in a more compact yet more accessible and dynamic state.

## Results

### HP1*β* segregates away from densely packed chromatin in mESCs

In most mouse cell types, the distribution in the nucleus of the three HP1 proteins (HP1 *α*, HP1*β* and HP1 *γ*) largely follows that of the DNA, being strongly enriched in heterochromatin in chromocentres. Generally, HP1 *α* is most faithfully associated with constitutive heterochromatin, while the HP1*β* and HP1 *γ* isoforms are also localised in euchromatin (Minc *et al*, 1999), (Dialynas *et al*, 2007), (Mattout *et al*, 2015). Indeed, HP1 *γ* has been suggested to participate in active transcription (Vakoc *et al*, 2005), (Mateescu *et al*, 2008). Notably, the pattern of HP1 localisation correlates with the cell’s differentiation state, with pluripotent ESCs showing the least accumulation of HP1 in chromocentres (Dialynas *et al*., 2007), (Bártová *et al*, 2008), (Mattout *et al*., 2015). Because HP1*β* and HP1 *γ* are found in both euchromatin and heterochromatin, we reasoned that studying their biophysical properties (and chromatin) in different contexts could help reveal the general principles underlying heterochromatin formation and organisation. To that end, we constructed an endogenous knock-in mES cell line, where we fused a HaloTag to the endogenous *Cbx1* gene allowing us to covalently modify the HP1*β* protein with organic dyes and study it in both fixed and live cells (Figure S1).

We first visualised the distribution of HP1*β* in naïve mouse embryonic stem cells (mESCs) using Airyscan imaging and we were surprised to find numerous (mean 12 per nucleus) generally small HP1*β* foci that were typically located in euchromatin (Figures 1A, B). These “DNA-poor/HP1*β*-rich” foci were not observed in either our earlier studies of HP1*β* in mouse L cells (Murzina *et al*., 1999) or in numerous other studies in cell types such as mouse embryonic fibroblasts (Maison *et al*, 2002) or human C127 cells (Dialynas *et al*., 2007). We also observed that heterochromatin did not always contain an increased density of HP1*β* (Figures 1A, B) – in particular, some regions that appeared to be chromocenters were notably depleted of HP1*β*. We termed these foci “DNA-rich/HP1*β*-poor”. Again, this was quite different from what we had observed previously in mouse L or NIH 3T3 cells where DNA dense regions always appeared to be enriched in HP1*β* (Thiru *et al*., 2004). We noted, however, that this lack of colocalisation was observed in earlier studies of mESCs grown on feeder cells in the presence of leukaemia inhibitory factor (LIF) (Dialynas *et al*., 2007), (Mattout *et al*., 2015). We wanted to understand how these disparities arise, and we wondered whether we could use this knowledge to better understand the mechanisms of HP1*β* focus formation and the function of HP1*β* in heterochromatin. We therefore set out to study the different foci in more detail.

**Figure 1.**
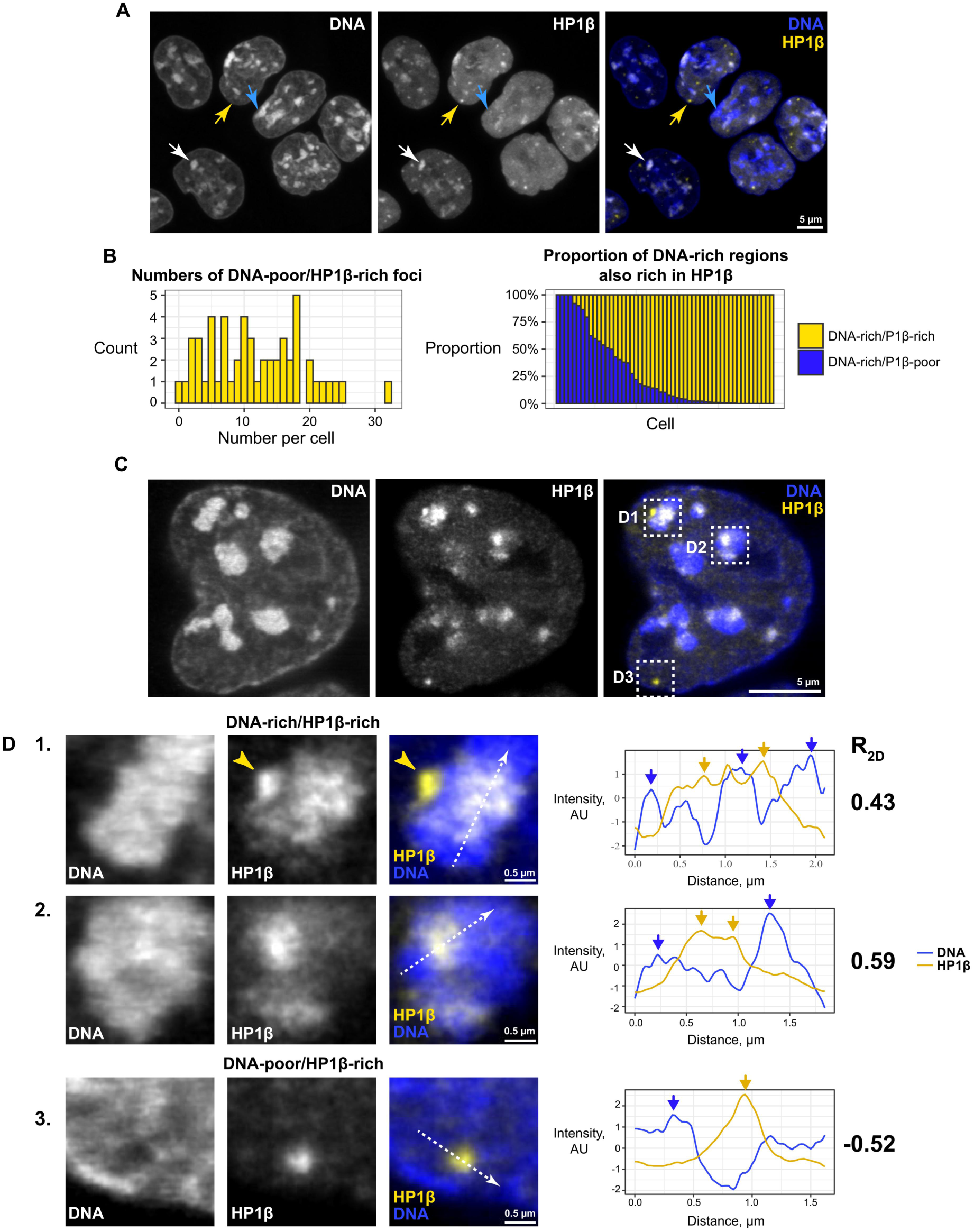
HP1*β* distribution differs from that of DNA in mES cells (see also Figure S2, Video S1). A. Airyscan confocal images of DNA (SiR-Hoechst) and HP1*β* (JF549-Halo) in mESCs (maximum projections of z stacks). Examples of DNA-poor/HP1*β*-rich (yellow arrow), DNA-rich/HP1*β*-poor (blue arrow) and canonical heterochromatic DNA-rich/HP1*β*-rich foci (white arrow) are indicated. B. Left, a histogram showing the number of DNA-poor/HP1*β*-rich foci per cell. Right, a quantification of the proportion of the DNA-rich/HP1*β*-poor and DNA-rich/HP1*β*-rich heterochromatic foci per cell (corrected by focus area). N = 53 cells from two independent replicates. C. 3D STED images of DNA (JF646-Hoechst) and HP1*β* (AF594-Halo) in mESCs (single slices). D. Left, close-ups of the areas annotated in C. In D1, yellow arrows indicate a DNA-poor/HP1*β*-rich focus associating with a larger DNA-rich/HP1*β*-rich area. Right, normalised line profiles along the lines shown in the images. The arrows highlight the anticorrelated peaks in the two channels. Rightmost, 2D Pearson’s correlation coefficient between the DNA and HP1*β* distribution within the foci shown. The segmentation procedure is illustrated in Figure S3A.

We noticed that even when HP1*β* and DNA were enriched in the same area, the shape of the focus often differed in the two images. We therefore asked whether HP1*β* and DNA within foci colocalise at high resolution using two colour 3D STED to obtain super-resolved images with fine optical sectioning. Previous studies had shown that HP1*α* is not uniformly distributed within foci (Erdel *et al*., 2020), (Novo *et al*, 2022) and we also found that this is the case for HP1*β* (Figure 1C). Surprisingly, however, we found that within “DNA-rich/HP1*β*-rich” foci, HP1*β* density was often locally anticorrelated with that of DNA (Figures 1C, 1D1, 1D2). We also observed that DNA-poor/HP1*β*-rich foci usually exist in pockets where the DNA density appears to be even lower than that in euchromatin (Figures 1C, 1D3). Interestingly, this suggested that the mechanism of concentration of HP1*β* in the DNA-rich/HP1*β*-rich and DNA-poor/HP1*β*-rich foci may be fundamentally similar. We also noted that the small DNA-poor/HP1*β*-rich foci often associate with a DNA-rich/HP1*β*-rich region (Figure 1D1) and we wondered whether this represents an example of the early stages of formation of a larger HP1-rich region. However, thus far we have not been able to observe any such merging of foci in live-cell time-lapse imaging of HP1*β* using either Airyscan, lattice light-sheet or eSPIM light-sheet microscopy (Video S1). In addition, we wondered whether the absence of HP1*β* in some DNA-rich foci was due to replacement of HP1*β* by HP1 *α*, but further studies showed that HP1 *α* was not present in foci which did not contain HP1*β* (Figure S2).

### HP1*β* binding in chromatin does not always correlate with histone H3K9me3

All HP1 proteins bind the histone H3K9me3 mark, which is thought to direct their distribution on chromatin (Bannister *et al*., 2001). We thus hypothesised that histone H3K9me3 levels might govern the unusual spatial pattern of HP1*β* that we observed. To investigate this, we performed three-colour Airyscan imaging to visualise DNA, HP1*β* and histone H3K9me3 (Figure 2A). Quantitative analysis of their distributions indicated that histone H3K9me3 correlates better with either HP1*β* or DNA density than HP1*β* and DNA density do with each other (Figure 2B). Surprisingly, we found that while histone H3K9me3 was often present in the DNA-rich/HP1*β*-rich foci, detailed examination of the images showed a notable lack of precise co-localisation of HP1*β* and histone H3K9me3 (Figures 2A, 2C5). In addition, while some DNA-poor/HP1*β*-rich puncta contained a lot of histone H3K9me3 (Figures 2A, 2C2), more typically the H3K9me3 signal appeared weak (Figures 2A, 2C1). The same was true for the DNA-rich/HP1*β*-poor foci where some were enriched for H3K9me3 (Figures 2A, 2C4), while others were not (Figures 2A, 2C3).

**Figure 2.**
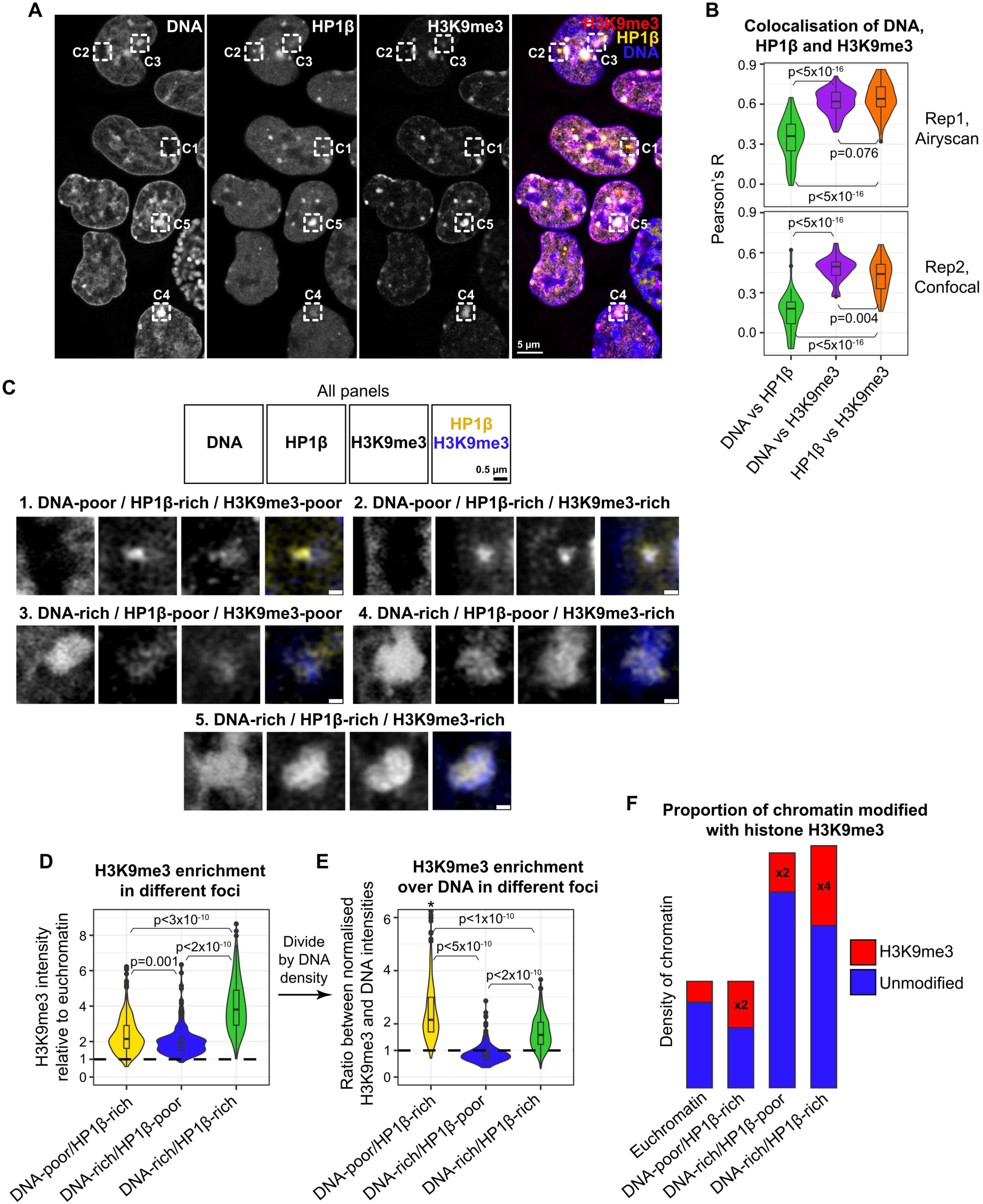
HP1*β* binding in chromatin does not always correlate with histone H3K9me3. A. Airyscan confocal images of DNA (Hoechst), HP1*β* (JF646-Halo) and H3K9me3 (immunofluorescence, Cy3) in mESCs (single slice). Examples of: DNA-poor/HP1*β*-rich foci with (C2) and without (C1) histone H3K9me3 enrichment; DNA-rich/HP1*β*-poor foci with (C4) or without histone H3K9me3 enrichment (C3); and canonical DNA-rich/HP1*β*-rich/histone H3K9me3-rich heterochromatin (C5) are shown. B. Quantification of DNA, HP1*β* and H3K9me3 colocalisation by Pearson’s correlation coefficient. Two biological replicates were imaged, using two different microscopes and different DNA dyes (either Hoechst or SPY505-DNA), and the results are thus shown separately. N = 69 and 44 cells per replicate, significance calculated using repeated-measures ANOVA with Greenhouse-Geisser correction and pairwise *p*-values calculated with paired Student’s t-test with Hochberg correction. C. Expanded views of the foci annotated in A (contrast adjusted). D, E. Quantification of histone H3K9me3 levels in the different kinds of foci. In D, the mean fluorescence intensity of each focus, divided by the mean signal in euchromatin in the corresponding cell, is plotted. In E, the ratio between the normalised histone H3K9me3 enrichment (from D) and the normalised DNA signal is shown. N > 180 foci per category, from 113 cells imaged in 2 biological replicates. *, 5 outlier values are beyond the plotted range. Significance was tested with Welch’s ANOVA with Games-Howell post-hoc. F. An illustration of chromatin density (represented by the height of the bar) and the proportion modified with histone H3K9me3 (denoted by the red colour, with the fold-change compared to euchromatin indicated) in different environments. Note that the absolute numbers of modified and unmodified nucleosomes are not known, only the relative differences between euchromatin and the different foci. The segmentation procedure is illustrated in Figure S3A.

Next, the different types of foci in the DNA and HP1*β* images were segmented out (Figure S3A), and the average histone H3K9me3 enrichment within the focus relative to euchromatin was quantified (Figure 2D). Nearly all foci had a score greater than 1, indicating some degree of H3K9me3 accumulation compared to euchromatin. However, in accordance with the qualitative observations described above, in the DNA-rich/HP1*β*-rich foci histone H3K9me3 levels were on average two-fold higher than in the other foci. The higher levels of histone H3K9me3 could either result from more extensive modification or the increased density of the chromatin, or both. To distinguish between these possibilities, we normalised the average histone H3K9me3 enrichment score of each focus by its mean enrichment in DNA density (Figure 2E). The DNA-rich/HP1*β*-poor foci generally displayed a ratio <1, revealing that histone H3K9me3 in these regions of densely packed DNA is depleted compared to euchromatin – perhaps suggesting that the methylation machinery has restricted access to these regions. In contrast, in both the DNA-rich/HP1*β*-rich and especially in the DNA-poor/HP1*β*-rich foci, the chromatin is typically more extensively modified with this mark compared to euchromatin (Figure 2F).

Next, we further investigated the patterns of histone H3K9me3 and either DNA (Figure 3A) or HP1*β* (Figure 3B) distribution at high resolution using two-colour 3D STED. As observed earlier, histone H3K9me3 was enriched in some of the DNA or HP1*β* foci, but not in others. DNA and H3K9me3 distribution within foci sometimes differed only in detail (Figure 3C1), but often the signals were locally anticorrelated (Figure 3C2). Similarly, when H3K9me3 was present in large heterogeneous HP1*β* foci (judging from their morphology likely DNA-rich/HP1*β*-rich foci), their distribution also usually differed in detail (Figure 3D1) but it was sometimes substantially different (Figure 3D2). Quantitative analysis confirmed that at high resolution histone H3K9me3 and HP1*β* colocalise better than H3K9me3 and DNA (Figure 3E), although this colocalisation is still rather variable. This suggests that local accumulation of HP1*β* does not depend on very high histone H3K9me3 levels, pointing to an additional mechanism facilitating the formation of HP1*β* foci. Consistently, when histone H3K9me3 was enriched in the small and dense DNA-poor/HP1*β*-rich foci, usually the two signals did not overlap exactly within the focus (Figure 3D3).

**Figure 3.**
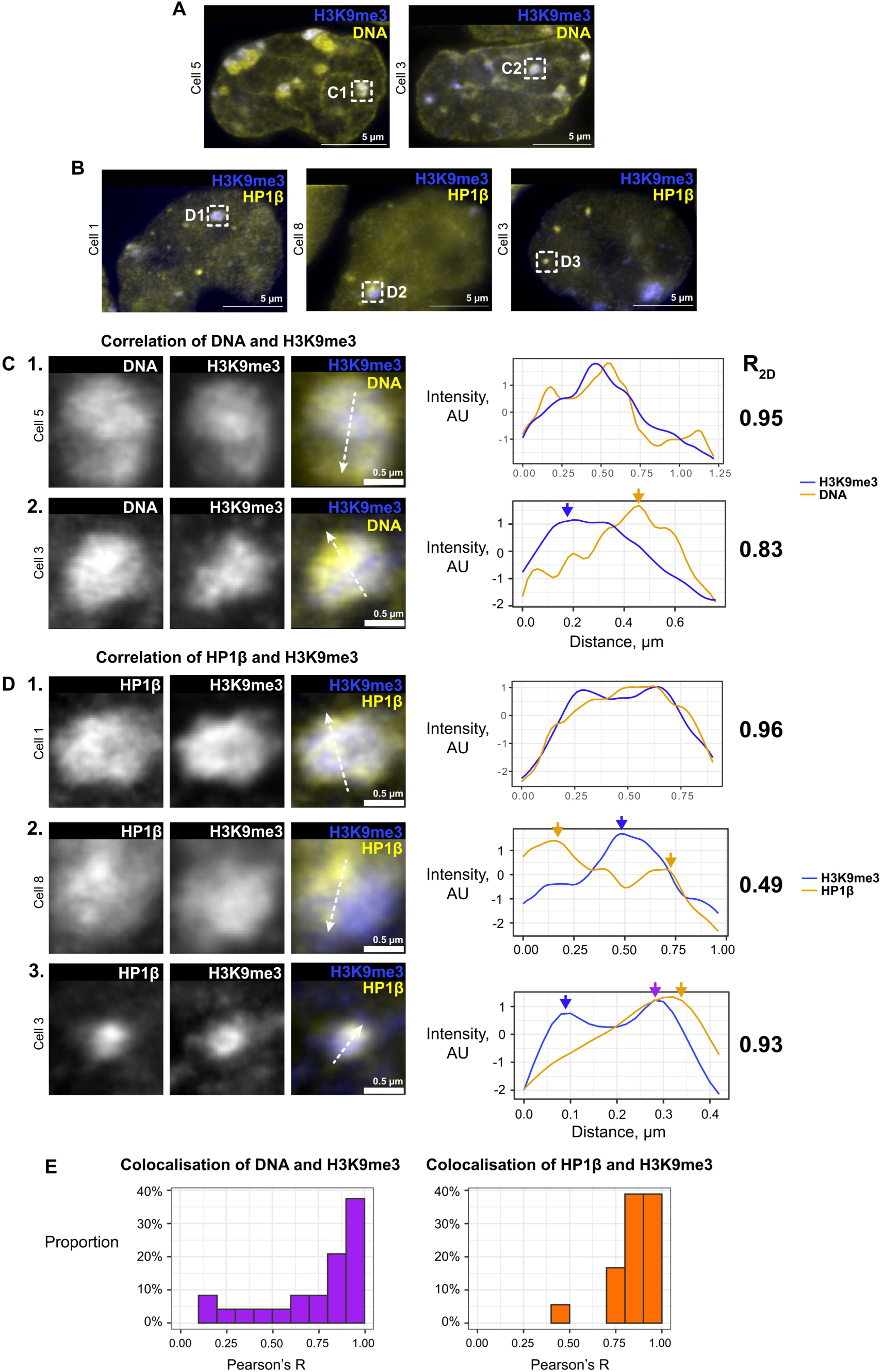
Colocalisation between H3K9me3 and HP1*β* on the nanoscale is variable, but more consistent than that between H3K9me3 and dense chromatin. A, B. 3D STED images of H3K9me3 (IF, AF594) and (A) DNA (JF646-Hoechst) or (B) HP1*β* (Halo-JF646) in mESC’s (single slices). C, D. Left, close-ups of foci annotated in A and B. Right, normalised line profiles along the lines shown in the images. The arrows highlight the correlated or anticorrelated peaks in the two channels. Rightmost, 2D Pearson’s correlation coefficient for the distribution between DNA and histone H3K9me3 (C), or HP1*β* and histone H3K9me3 (D) within the foci shown. E. 2D Pearson’s correlation coefficient for the distribution between DNA and histone H3K9me3 within foci (Left; 24 foci in 6 cells) or HP1*β* and H3K9me3 distribution within large foci likely to be heterochromatic (Right; 18 foci in 8 cells). The segmentation procedure is illustrated in Figure S3B.

### HP1*β* is concentrated in foci via a combination of stable chromatin binding and retention of mobile protein

Next, we set out to investigate the biophysical features of the different types of foci. Broadly speaking, HP1*β* could be concentrated locally via two different mechanisms: either *1)* canonical stable binding to chromatin, e.g. via chromodomain-histone H3K9me3 or shadow chromodomain interactions with either other protein complexes or the histone H3-*α* N helix (Bannister *et al*., 2001), (Murzina *et al*., 1999), (Richart *et al*, 2012); or *2)* via formation of a liquid phase-separated condensate through a multitude of weak interchangeable interactions (Larson *et al*., 2017), (Strom *et al*., 2017). Assuming that diffusion is unimpeded and that molecules can rapidly leave the foci upon dissociating from chromatin, the former model predicts that any local increase in HP1*β* concentration is caused by a higher chromatin-bound fraction of the protein, while the concentration of freely diffusing HP1*β* should remain equilibrated with the rest of the nucleoplasm. On the other hand, if weak multivalent HP1*β*-HP1*β*, or HP1*β*-chromatin interactions are responsible for condensation, some mobile chromatin-unbound HP1*β* is expected to be retained within the foci.

To test these predictions in the context of the foci observed in mESCs, the diffusion of single HP1*β* molecules in live cells was tracked and analysed. HP1*β* is known to be less mobile in heterochromatin than in euchromatin (Cheutin *et al*., 2003), (Festenstein *et al*., 2003), (Schmiedeberg *et al*, 2004), which could result from either a higher chromatin-bound fraction and/or slower diffusion of the unbound molecules. Because the diffusion coefficients of molecules that are either bound or not bound to chromatin differ by orders of magnitude (Lippincott-Schwartz *et al*, 2001), (Akhtar & Gasser, 2007), single molecule tracking allows one to estimate the bound fractions of each species and infer their apparent diffusion coefficients. We imaged mESCs for over two minutes in two colours – HP1*β* was tracked with a frame rate of 286 Hz (exposure time 3.5 ms) in one channel, and DNA was imaged in the other. This allowed us to find and segment out the three types of foci based on their DNA and HP1*β* enrichment – the remainder of the cell was labelled as euchromatin (Figure S4). The molecular trajectories were then classified based on their location. Importantly, because the segmentation procedure places geometric constraints on the types of trajectories that can “fit” into the various foci, the estimation of diffusion parameters may be biased. Therefore, as a control, the shapes of the segmented foci were randomly displaced into euchromatin within the same nucleus, and the diffusion parameters were also estimated for the trajectories in the resulting “pseudo foci” for comparison (Figures S4B,C).

We applied the new SA-SPT method (Heckert *et al*, 2022) to analyse 100,000 trajectories and recover the spectrum of apparent diffusion coefficients that the HP1*β* molecules display (Figure 4A). Firstly, a population of essentially stationary molecules was detected, which we attribute to chromatin-bound HP1*β* and whose diffusion coefficient is not measurable at this frame rate. In addition, at least two distinct mobile states could also be observed, with apparent diffusion coefficients around 1 μm^2^/s and 2.5 μm^2^/s (Figure 4A). As a control, the fluorescent dye was immobilised on a coverslip. This sample showed a spectrum that nearly completely consisted of immobile molecules, while the “background” probability density was very low, demonstrating that the sample size is sufficient to clearly distinguish different diffusive states. Thus, the “background” density in the live-cell data (e.g. around D = 0.1 μm^2^/s) is likely not a consequence of a lack of sensitivity of the method but rather reflects underlying molecular processes such as fast transient binding of HP1*β* to chromatin. Both curves rapidly decayed to zero around D = 3.0-3.5 μm^2^/s, indicating the upper detection limit of the technique.

**Figure 4.**
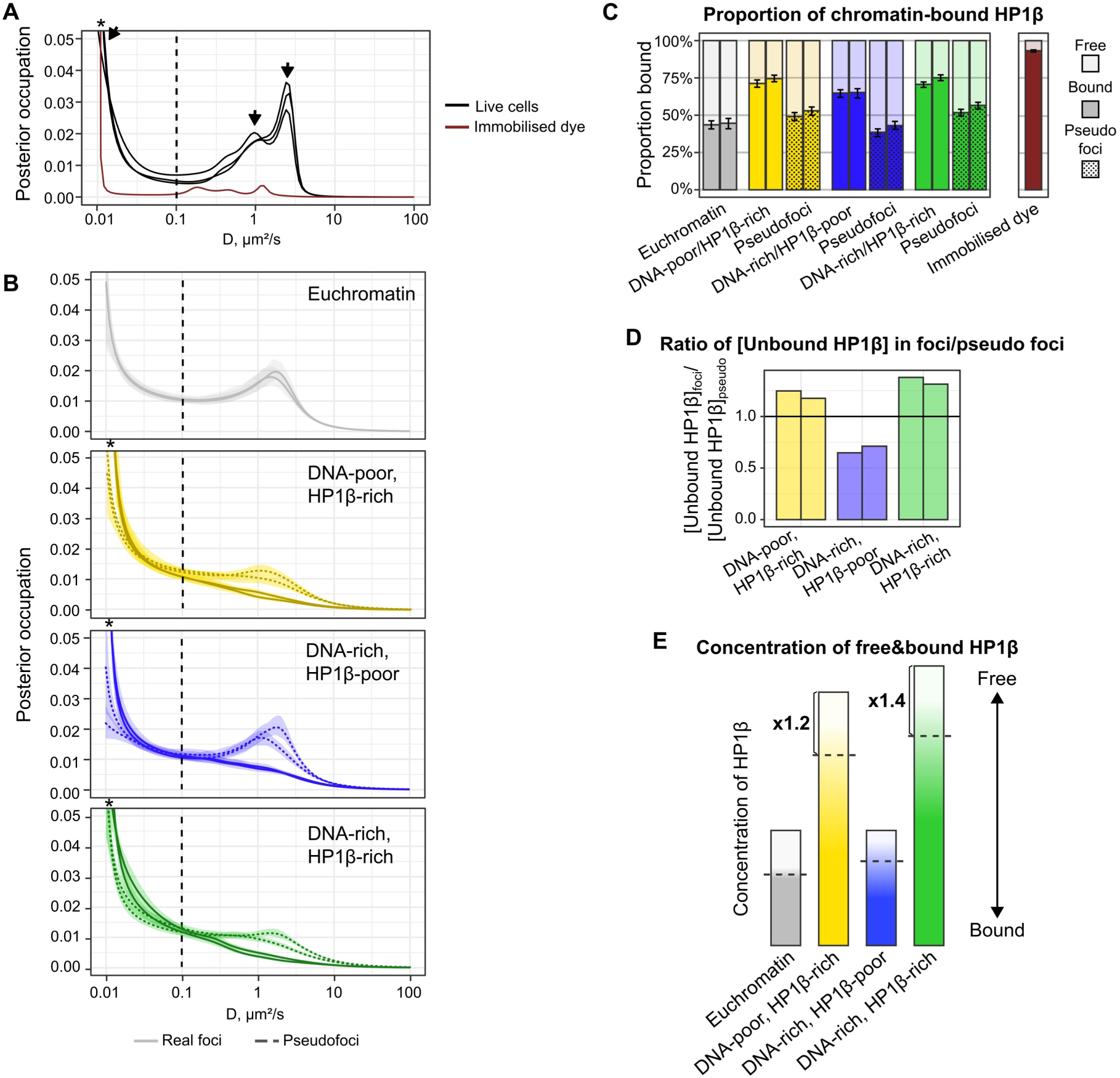
HP1*β* is concentrated in foci via a combination of stable chromatin binding and retention of mobile protein (see also Figures S4 and S5) A. Spectrum of apparent diffusion coefficients for HP1*β* in live cells (3 biological replicates, 11-21 cells each) as well as for Halo-JF639b dye immobilised on a coverslip (1 replicate), calculated from 100,000 trajectories per sample using SA-SPT. The arrows indicate the three populations of HP1*β*. B. Apparent diffusion coefficient distributions for HP1*β* in euchromatin and different nuclear foci and their corresponding control pseudo foci. 5,000 and 3,100 trajectories were subsampled in each environment in the two biological replicates, and the 95% confidence interval was generated by bootstrapping each 100 times. In A and B, the *’s show that the spectrum extends beyond the plot area, and the vertical dashed lines indicate the (arbitrary) threshold between the chromatin-bound and diffusing molecules that we used in the subsequent analysis. C. The proportion of chromatin-bound, i.e. immobile, HP1*β* molecules in different nuclear environments, calculated by integrating the SA-SPT posterior occupation density above and below the threshold (two replicates). Error bars indicate the 95% confidence interval, generated by bootstrapping 100 times. N = 20,000 trajectories for the immobilised dye control. D. The ratio of the concentrations of unbound HP1*β* within real foci and their corresponding pseudo foci controls, i.e. euchromatin (two replicates). E. An illustration of the concentration (represented by the height of the bar) and mobility (denoted by the colour) of HP1*β* in different environments. The gradual change in colour indicates that in reality, a range of diffusive states exist. To distinguish “chromatin-bound” from “freely diffusing” HP1*β* we binarised the spectrum using a threshold value of 0.1 μm^2^/s (dashed lines). The fold-change in concentration of the “free” molecules compared to euchromatin is shown.

Next, HP1*β* diffusion was analysed separately in the different foci (Figure 4B). To accurately compare the diffusion parameters, the data were subsampled to give equally-sized datasets – reducing the dataset size to the number of trajectories in the rarest foci (plots produced from complete datasets are shown in Figure S5). This reduced amount of data did lead to less well-defined populations compared to Figure 4A. Strikingly, all types of foci showed a marked decrease in the proportion of mobile molecules compared to either their pseudo foci controls or euchromatin (Figures 4B, S5). To quantify this, a threshold value of D = 0.1 μm^2^/s based on the well-defined distribution from Figure 4A was selected to estimate the relative numbers of mobile and immobile molecules. The proportion of chromatin-bound HP1*β* in euchromatin was about 45%, but in DNA-rich/HP1*β*-poor foci it increased to roughly 65%, and in both DNA-poor/HP1*β*-rich and DNA-rich/HP1*β*-rich foci to 70-75% (Figure 4C). Furthermore, the broad peak corresponding to mobile HP1*β* observed in euchromatin (likely to be multiple overlapping peaks, see Figure 4A) essentially vanished in all types of foci, leaving a “smear” without any discrete states. We suggest that this is because nearly all the HP1*β* molecules are engaged in transient interactions. Indeed, such “smearing” is observed in simulations where the time scale of state transitions or binding-unbinding is comparable to that of diffusion (Heckert *et al*., 2022).

The liquid-liquid phase separation (LLPS) model predicts that freely diffusing or weakly bound molecules should be retained within the condensate. In other words, the concentration of such molecules within the focus is expected to be higher than outside. To test this prediction, we calculated the enrichment of chromatin-unbound HP1*β* within foci compared to their pseudo foci (Figure 4D), using the fractions shown in Figure 4C and the local density of HP1*β* from the SPT data. Indeed, we found a 1.2-1.4-fold enrichment in chromatin-unbound HP1*β* in both the DNA-rich/HP1*β*-rich and DNA-poor/HP1*β*-rich foci compared to euchromatin (Figure 4D). Thus, increased HP1*β* concentration in these foci is caused not only by an increase in stable chromatin binding, but also by the retention of mobile HP1*β* molecules. In contrast, the concentration of freely diffusing HP1*β* molecules in DNA-rich/HP1*β*-poor heterochromatic foci was found to be about 30% lower than in euchromatin (Figure 4D). It thus appears that either the high chromatin density sterically hinders, or some other factor limits the diffusion of mobile HP1*β* into these foci.

We next investigated whether the increase in the proportion of chromatin-bound HP1*β* in the different types of foci could arise from a higher density of available histone H3K9me3 binding sites, assuming a simple non-variable bimolecular reaction mechanism:

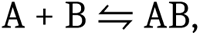

where A is [HP1*β*] and B its [histone H3K9me3 binding sites on chromatin]. (This assumption is expected to be valid when the concentration of histone H3K9me3 binding sites is low – at higher concentrations the dimeric HP1*β* protein can bind through both its chromodomains and exhibit more complex kinetics.) We found that this kinetic scheme could explain the data for the DNA-rich/HP1*β*-poor and the DNA-rich/HP1*β*-rich foci. However, the equation Kd(euchromatin) = Kd(DNA-poor/HP1*β*-rich foci) has no solution when we take the concentration of B as proportional to histone H3K9me3 density. The concentration of histone H3K9me3 binding sites is expected to be low in DNA-poor/HP1*β*-rich foci – see the assumption above and the Supplementary Note. Thus, to explain the very high (70-75%) proportion of chromatin-bound HP1*β* within DNA-poor/HP1*β*-rich foci, a change in the mechanism of binding of HP1*β* to chromatin must be involved, e.g. due to cooperative effects and/or LLPS.

In summary, in both DNA-poor/HP1*β*-rich and DNA-rich/HP1*β*-rich foci, HP1*β* appears to be locally concentrated through a combination of increased HP1*β* binding to chromatin and the retention of freely diffusing molecules (Figure 4E). The disappearance of the distinct mobile population in the spectra (Figures 4B, S5) is consistent with most of the “free” HP1*β* being involved in transient interactions, which might also facilitate stable binding to chromatin. Overall, these results are consistent with the formation of a dynamic liquid phase-separated condensate forming around stably chromatin-bound HP1*β* molecules. In contrast, the DNA-rich/HP1*β*-poor foci display an enrichment in chromatin-bound HP1*β* apparently because chromatin compaction provides more binding sites for the protein. However, they are depleted of mobile HP1*β*, potentially due to steric hindrance.

### Both the density and the abundance of HP1*β* foci decrease upon reduction in HP1*β* concentration

In addition to their retention of unbound protein, another distinguishing feature of liquid phase-separated systems is their response to changes in the concentration of their constituents (Banani *et al*., 2017), (McSwiggen *et al*, 2019). In a simple LLPS model, an increase in the solute (HP1*β*) concentration leads to an increase in the volume fraction of the dense phase (foci), but not the concentration of the solute in each phase (“concentration buffering”). In contrast, if the recruitment of HP1*β* to foci is governed by specific stable macromolecular interactions with appropriately modified chromatin, one would expect the concentration of HP1*β* both inside and outside of foci to rise upon an increase in total concentration, while the volume fractions of the two environments should remain the same (“size buffering”).

We therefore compared wild-type mESCs with cells where one allele of *Cbx1* was knocked out (*Cbx1*^+/-^), which express two-fold lower levels of the HP1*β* protein (Figure S1D). The diffuse euchromatic HP1*β* labelling in the *Cbx1*^+/-^ mutant cells was weaker, with fluorescence intensity values approximately two times lower than in the wild-type. Furthermore, this was also the case in both the DNA-rich/HP1*β*-rich and DNA-poor/HP1*β*-rich foci, while their relative intensity with respect to euchromatin did not change (although statistically significant, the changes are <5%, see Figures 5A, 5B). However, the DNA-poor/HP1*β*-rich foci were less numerous in the low-expressing cells (Figures 5A, 5C), while the areas covered by the DNA-rich/HP1*β*-rich or DNA-rich/HP1*β*-poor heterochromatin remained constant (data not shown). The decrease in HP1*β* concentration in both the DNA-rich/HP1*β*-rich and DNA-poor/HP1*β*-rich foci upon a reduction in the total amount of HP1*β* protein indicates clearly that the foci are not simple liquid phase separated condensates, which is also consistent with the high proportion of stably chromatin-bound HP1*β* (70-75%). On the other hand, the reduced number of DNA-poor/HP1*β*-rich foci in the *Cbx1*^+/-^ mutant cells suggests that LLPS does play a role in the formation of these foci and potentially reflects a lower probability of nucleation of phase-separated condensates at lower HP1*β* concentrations.

**Figure 5.**
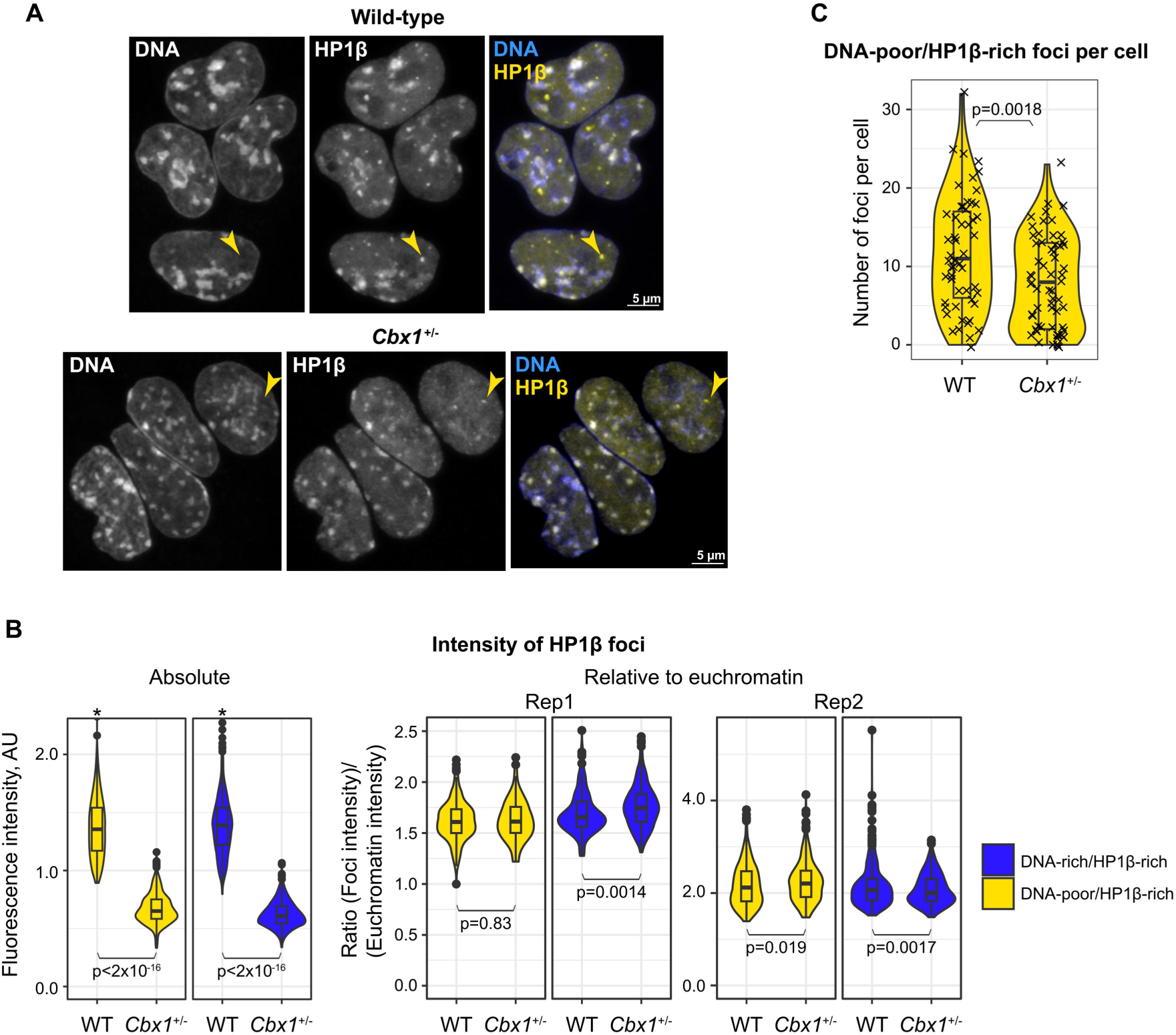
Both the density and the abundance of HP1*β* foci decrease upon reduction in HP1*β* concentration. A. Airyscan confocal images of DNA (SiR-Hoechst) and HP1*β* (JF549-Halo) in wild-type and mutant *Cbx1^+/-^* cells expressing two-fold less HP1*β* (maximum projections of stacks). Examples of DNA-poor/HP1*β*-rich foci are shown with yellow arrows. (The intensities in the images were adjusted for presentation and are not comparable between the two panels.) B. Quantification of the HP1*β* intensity (left, absolute values; right, relative to the intensity of euchromatin) of DNA-poor/HP1*β*-rich or DNA-rich/HP1*β*-rich foci in wild-type and *Cbx1^+/-^* mESCs. N = 628 and 485 foci in 53 and 61 cells, respectively, from two biological repeats. Because the intensities of the repeats were different, the fluorescent intensities were internally normalised to arbitrary units (left), and the ratios are shown in separate panels (right). The *’s show 3 and 2 outliers lying beyond the plotted area. C. Quantification of the number of DNA-poor/HP1*β*-rich foci per cell in wild-type and *Cbx1*^+/-^ mESCs. N = 53 and 61 cells, two biological repeats. Statistical significance was determined by two-tailed Student’s or Welch’s t-tests.

### HP1*β*-associated heterochromatin is denser, but more dynamic

Recent studies have shown that the dynamics of chromatin may play a significant role in gene regulation and other chromatin-associated processes (Gu *et al*, 2018), (Germier *et al*, 2017), (Oshidari *et al*, 2020). Heterochromatin is generally less dynamic than euchromatin (Nozaki *et al*, 2017), (Lerner *et al*, 2020), which likely contributes to its repressive state by limiting the contact probability between different regions of the genome. Furthermore, the dense packing of DNA in heterochromatin could also potentially restrict the access of large protein complexes, leading to transcriptional repression. Since HP1 can cross-link chromatin fibres (Hiragami-Hamada *et al*., 2016), it is conceivable that a large amount of chromatin-bound HP1 might lead to a lower chromatin mobility. On the other hand, this effect might be dampened by the dynamic nature of HP1 interactions with chromatin. We thus set out to measure chromatin mobility in the different types of foci to investigate the separate and combined effects of HP1*β* and chromatin condensation.

To this end, we employed spinning-disk confocal imaging of DNA and HP1*β* in combination with Hi-D, an analysis method that estimates local diffusion parameters using optical flow (Shaban & Seeber, 2020). One hundred and fifty images with a 200 ms exposure were acquired, summing up to 30 s of imaging per nucleus. Compared to fast-exposure SPT, the much longer exposure time employed in this experiment allowed us to observe and quantify the slow movement of chromatin. An anomalous diffusion model was used to fit the data, and a directional movement component was included to account for large-scale motions. Figure 6A shows the distributions of diffusion coefficients per pixel in different environments. Remarkably, chromatin diffusion was only markedly different from that in euchromatin in a proportion of DNA-rich/HP1*β*-poor foci, where it was notably lower. In contrast, chromatin mobility was very similar to that in euchromatin in both the DNA-rich/HP1*β*-rich and DNA-poor/HP1*β*-rich foci, a result that is consistent with previous work showing that HP1*β* is depleted from very low mobility chromatin (Lerner *et al*., 2020).

**Figure 6.**
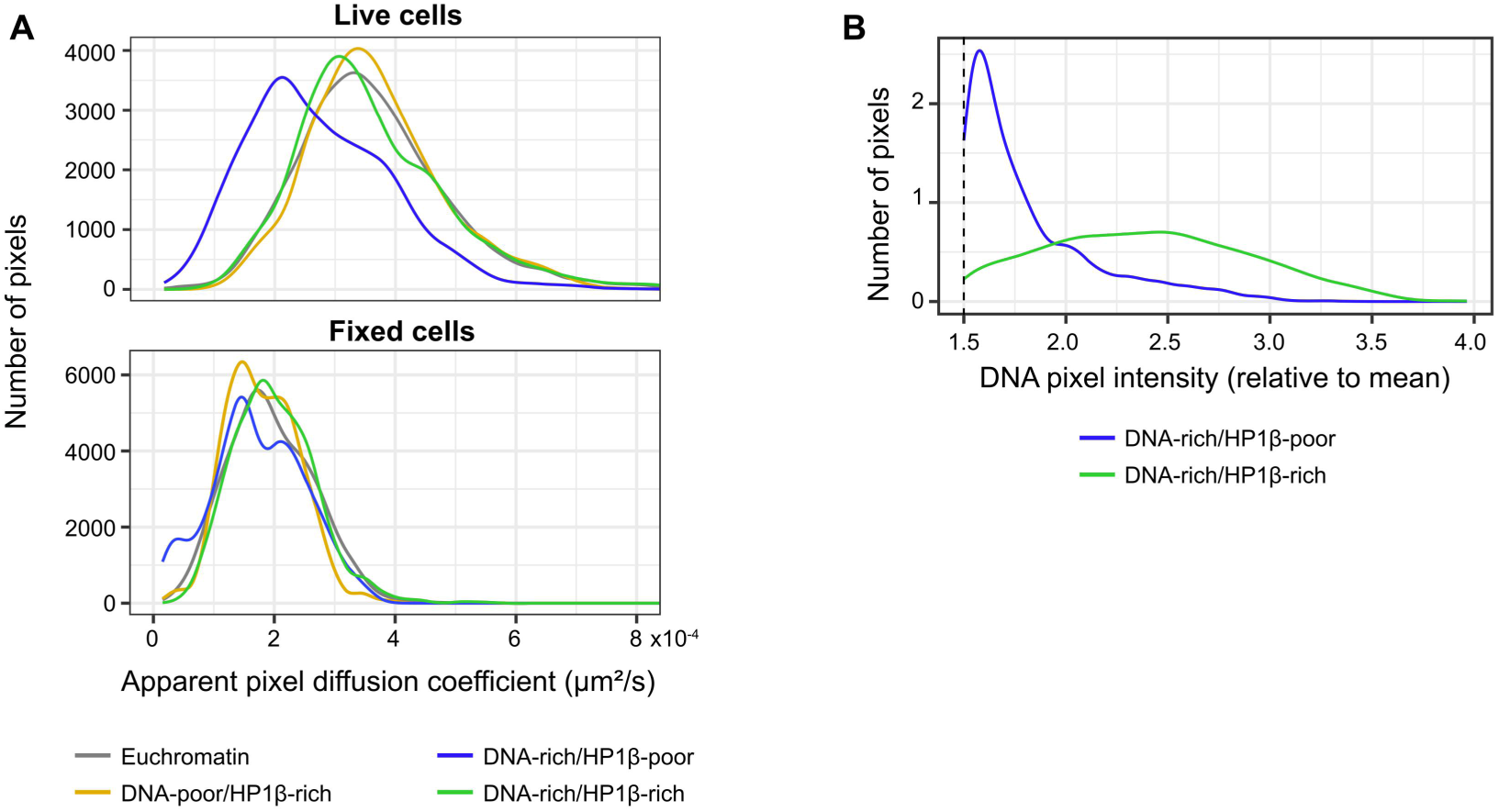
Hi-D indicates that HP1*β*-associated heterochromatin is denser, but more dynamic. A. Distribution of diffusion coefficients, estimated per pixel by Hi-D. An equal number of pixels was subsampled from each cell and each environment. Top, the data for live cells. Two biological replicates (15 and 19 cells), each of which showed a similar trend, were pooled. Bottom, fixed-cell control, single replicate (11 cells). B. Distributions of DNA signal intensity per pixel, relative to the mean within that cell, in DNA-rich/HP1*β*-rich and DNA-rich/HP1*β*-poor heterochromatin. The dashed line indicates the threshold for defining heterochromatin based on DNA signal intensity.

To test whether the decreased chromatin mobility within DNA-rich/HP1*β*-poor compared to DNA-rich/HP1*β*-rich foci is associated with altered chromatin compaction, the intensity of the DNA signal in these foci was compared (Figure 6B). This analysis indicated that DNA-rich/HP1*β*-poor foci have a lower density of DNA, i.e. chromatin is less compacted than that in the DNA-rich/HP1*β*-rich foci. Thus, HP1*β* in DNA-rich/HP1*β*-rich heterochromatin is associated with tightly packed, but mobile chromatin.

### Canonical heterochromatin forms upon exit from naïve pluripotency

Previous work has shown that chromosome structure changes considerably when ES cells exit from naïve pluripotency (Novo *et al*, 2016), (Lando *et al*, 2024). Furthermore, the levels of HP1*β* are thought to be increased in pluripotent cells compared to more differentiated cells, and HP1*β* was found to be crucial to both the maintenance of pluripotency and the proper differentiation of ESCs (Mattout *et al*., 2015). We therefore investigated HP1*β* distribution during the onset of mESC differentiation in an *in vitro* model that recapitulates the very early stages of mouse development (Kalkan *et al*, 2017). In this model, differentiation is initiated by removing two small molecule inhibitors and LIF (2i/LIF) from the culture. In previous work we found that, in addition to a change in the physical “softness” of the nuclei 24 hrs after removal of 2i/LIF (Pagliara *et al*, 2014), as cells enter the formative state there is a striking increase in long-range contacts, chromosome intermingling and the formation of multiway chromatin hubs (Lando *et al*., 2024). These changes in chromosome structure are mostly reversed by 48 hrs, and we wondered whether this formative state transition involves a breakdown in heterochromatin structure and its subsequent reestablishment.

We first compared the levels of HP1*β* at these early stages of differentiation. We found that HP1*β* protein levels increase by about 50-80% 24 hrs after removal of 2i/LIF, and that this is at least partially due to transcriptional upregulation of the gene (Figure 7A, Figures S6A, S6B). Unexpectedly, at the same time, global levels of the histone H3K9me3 modification decreased – as assessed by histone H3K9me3 levels on major satellite (Figure 7B) and other repeats (Figure S6C).

**Figure 7.**
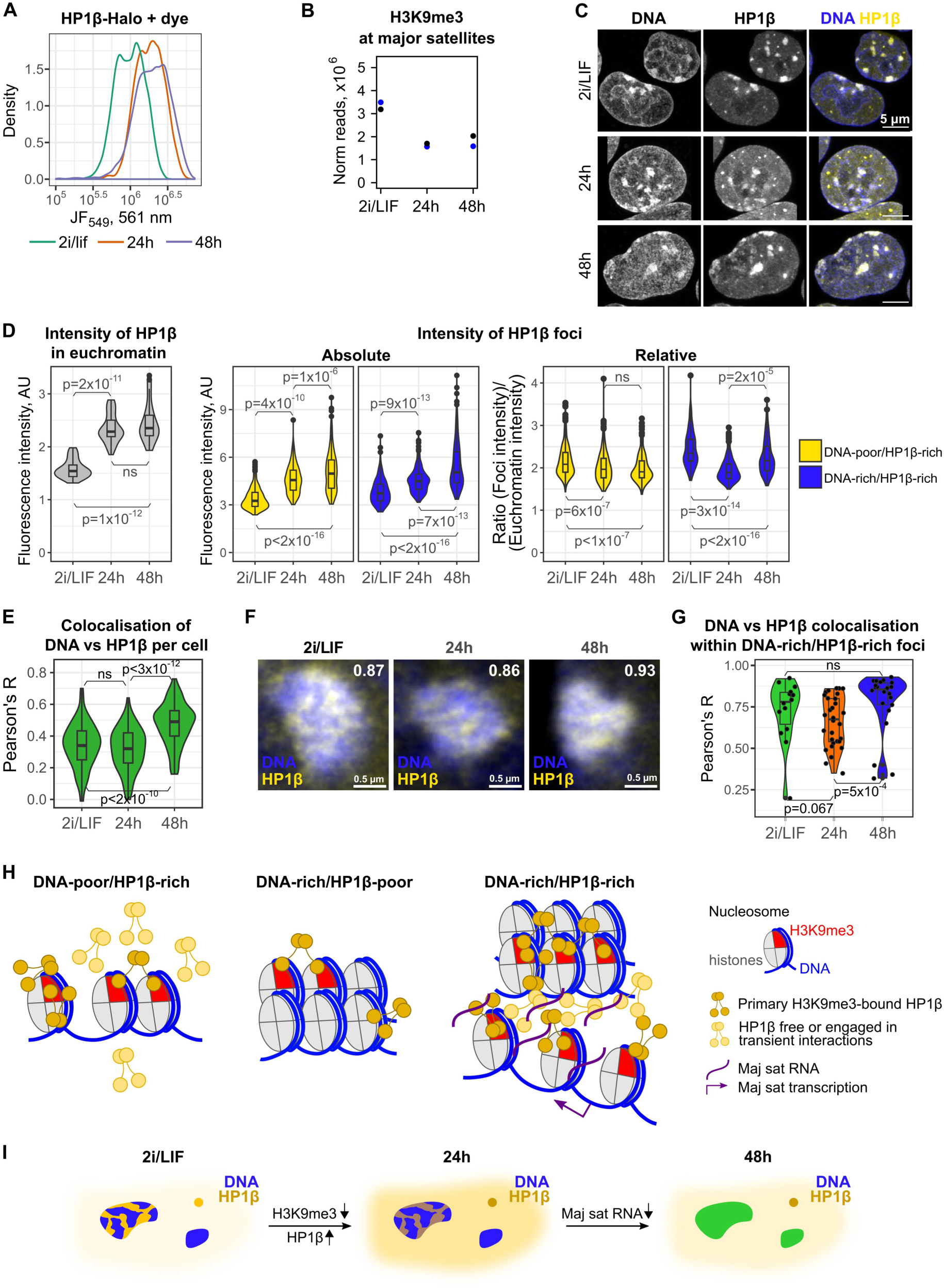
Canonical heterochromatin forms upon exit from naïve pluripotency (see also Figures S6 and S7). A. Histograms of the level of fluorescence intensity in single cells in different conditions, measured using flow cytometry after labelling HP1*β*-Halo with an organic dye Halo-JF549. B. Levels of histone H3K9me3 at major satellite repeats, measured using quantitative ChIP-seq (2 replicates). C. Airyscan confocal images of HP1*β* (Halo-JF646) and DNA (Hoechst) in mESCs in 2i/LIF and either 24 or 48 hrs after the onset of differentiation (maximum projections of z-stacks; see more examples in Figure S7A). (The displayed intensities are adjusted for better visibility, and are thus not comparable between images.) D. Quantification of HP1*β* intensity. Left, mean intensity in euchromatin per cell (N = 33, 36 and 27 cells); significance tested with one-way ANOVA with Tukey post-hoc. Middle and right, intensity of DNA-rich/HP1*β*-rich and DNA-poor/HP1*β*-rich foci (middle, absolute values; right, relative to the intensity of euchromatin; >200 foci in each group). Statistical significance tested with ANOVA or Welch’s ANOVA with Tukey or Games-Howell post-hocs. E. Colocalisation analysis during the differentiation time course, performed by calculating Pearson’s correlation coefficient between pixel intensities in the DNA and HP1*β* channels on a per-nucleus basis. Statistical testing with one-way ANOVA with Tukey’s post-hoc. F. 3D STED images of DNA-rich/HP1*β*-rich foci (DNA, JF646-Hoechst; HP1*β*, Halo-JF646) in mESC’s, and the corresponding 2D Pearson’s correlation coefficients between the DNA and HP1*β* distributions (see full images in Figure S7B). G. Comparison of 2D Pearson correlation coefficients for all DNA-rich/HP1*β*-rich foci in the STED images. Statistical testing with Kruskal-Wallis test (p=0.004) with Dunn post-hoc. H. A model for the organisation of the heterochromatic foci in mES cells. The DNA-poor/HP1*β*-rich foci (left) contain heavily H3K9me3-modified chromatin, to which HP1*β* binds. This seeds the formation of a condensate, where HP1*β* is engaged in a variety of stable and transient interactions with chromatin and other components and displays a range of diffusive behaviours. The DNA-rich/HP1*β*-poor foci (middle) contain cross-linked and immobile chromatin. HP1*β* is predominantly bound to the H3K9me3 mark within these regions, while the mobile protein is excluded. The DNA-rich/HP1*β*-rich foci (right) contain regions of high chromatin density (top part) and HP1*β* pockets (bottom part). We hypothesise that the latter is associated with major satellite (or other) RNA. I. A model for heterochromatin reorganisation during early differentiation. In naïve mESCs (2i/LIF conditions), DNA-rich/HP1*β*-poor, DNA-poor/HP1*β*-rich and DNA-rich/HP1*β*-rich foci are present, with dense chromatin and HP1*β* being segregated in the latter. 24 hrs after the onset of differentiation, decreased histone H3K9me3 levels and increased HP1*β* expression result in a weaker enrichment of HP1*β* in DNA-rich/HP1*β*-rich foci compared to euchromatin. At 48 hrs, the enrichment is restored, and HP1*β* and DNA densities become more correlated (shown with a green colour), potentially due to a decrease in major satellite (or other) RNA transcription.

To investigate the combined effects of increased HP1*β* levels and decreased numbers of histone H3K9me3 binding sites on heterochromatin structure, we studied the distribution of HP1*β* and DNA in 2i/LIF and after 24 and 48 hrs of differentiation using Airyscan imaging (Figures 7C, S7A). The intensity of HP1*β* in euchromatin as well as that in both the DNA-poor/HP1*β*-rich and DNA-rich/HP1*β*-rich foci increased at 24 and 48 hrs (Figures 7C, 7D, S7A). However, the ratio of HP1*β* levels in DNA-rich/HP1*β*-rich foci to that in euchromatin actually decreased at 24 hrs, whilst that in the DNA-poor/HP1*β*-rich foci did not change (Figures 7C, 7D, S7A). In contrast to the results obtained for the *Cbx1^+/-^* mutant (Figure 5C), the number of DNA-poor/HP1*β*-rich foci did not change despite the elevated HP1*β* protein concentration (data not shown), suggesting that the increased levels of HP1*β* may be countered by the loss in histone H3K9me3 binding sites. Surprisingly, despite the absence of major changes in either HP1*β* or H3K9me3 levels between 24 hr and 48 hr cells (Figures 7A, 7B), the DNA and HP1*β* images became much more similar at 48 hrs – an observation that was confirmed by colocalisation analysis (Figure 7E).

To investigate the changes in the detailed structure of foci, we carried out two-colour 3D STED imaging of HP1*β* and DNA at the three stages (Figure 7F, Figure S7B). At the ultrastructural level, it appears that the colocalisation of HP1*β* and DNA within DNA-rich/HP1*β*-rich foci decreases at 24 hrs after 2i/LIF removal, and then increases at 48 hrs (Figures 7F, 7G, S7B), although the statistical significance of the quantitative differences is low due to the limited throughput of STED imaging (Figure 7G).

In summary, HP1*β*-mediated heterochromatin structure appears to be partially broken down as cells enter formative pluripotency 24 hrs after 2i/LIF withdrawal, and subsequently it is reestablished in a more canonical form. Whilst the relative changes in HP1*β*/H3K9me3 levels are likely to be in part responsible for the redistribution of HP1*β* during the first 24 hrs of differentiation, some other process must underlie the changes that occur between 24 hrs and 48 hrs – because neither the HP1*β* nor the H3K9me3 levels change between 24 hrs and 48 hrs. We hypothesise that some additional factor that is responsible for segregating HP1*β* away from dense chromatin is produced during early differentiation that later disappears.

## Discussion

In this study we discovered that both DNA-rich/HP1*β*-poor and DNA-poor/HP1*β*-rich foci exist alongside canonical DNA-rich/HP1*β*-rich heterochromatin in naïve pluripotent mouse ES cells (Figure 1A). Furthermore, even within the canonical DNA-rich/HP1*β*-rich heterochromatic foci, regions of high HP1*β* and DNA are often segregated away from each other (Figures 1C, 1D), which stimulated us to reexamine how heterochromatin forms. We found that the levels of H3K9me3 in foci are extremely variable, and do not always correlate with those of HP1*β*. Nevertheless, there is always some enrichment of H3K9me3 compared to euchromatin (Figure 2). We thus propose that some local accumulation of H3K9me3 is crucial for seeding an HP1*β*-rich focus by providing a platform for stable chromatin binding. Subsequently, H3K9me3 levels in foci might increase in a maturation step that stabilises HP1*β* association, which may be important for reliable heritability of HP1*β*-mediated heterochromatin through cell division. This process could involve shadow chromodomain-mediated recruitment of Lysine-9 histone H3K9 methyltransferases such as SetDB1 (see e.g. (Schultz *et al*., 2002), (Loyola *et al*, 2009)). Alternatively, HP1*β* may be recruited and locally concentrated *via* histone H3K9me3-independent mechanisms, e.g. *via* histone H1.4 (Daujat *et al*, 2005).

Single-particle tracking indicated that all three types of HP1*β* foci contain a higher proportion of chromatin-bound HP1*β* compared to euchromatin (Figures 4B, 4C), consistent with their increased local histone H3K9me3 concentration (Figure 2D). However, the DNA-poor/HP1*β*-rich and DNA-rich/HP1*β*-rich foci also retain freely diffusing HP1*β* molecules, which we suggest could be engaged in transient interactions forming a liquid phase-separated condensate (Figure 4D). These freely diffusing HP1*β* molecules are depleted from DNA-rich/HP1*β*-poor foci (Figure 4D). Altering the total amount of HP1*β* protein in cells led to a proportional change in the concentration of HP1*β* within foci, suggesting that stable chromatin binding is the major factor in their formation and maintenance (Figures 5A, 5B). This is consistent with the fact that HP1*β* foci nearly always disperse during mitosis, when HP1 proteins are displaced from chromatin by histone H3S10 phosphorylation (Minc *et al*., 1999), (Murzina *et al*., 1999), (Fischle *et al*, 2005). However, the number of DNA-poor/HP1*β*-rich puncta per cell was also affected (Figure 5C), suggesting that cooperative binding and/or LLPS may also play a significant role in the formation of HP1*β* foci. Thus, the DNA-poor/HP1*β*-rich foci display features that are consistent with a model combining stable chromatin binding and liquid-liquid phase separation via transient interactions.

What are the “DNA-poor/HP1b-rich” foci, and what is their function? HP1 proteins are known to localise to promyelocytic leukaemia (PML) nuclear bodies via their interaction with SP100 (Seeler *et al*, 1998) and the size (∼0.1-0.2 μm in diameter, Figure 1D) and morphology of the DNA-poor/HP1b-rich foci does resemble that of HP1-containing PML bodies found in HeLa cells (∼0.2-1.0 μm in diameter, Hayakawa et al 2003). PML bodies have also been reported in ESCs, and the PML protein appears to play a role in pluripotency maintenance (Hadjimichael *et al*, 2017), (Tessier *et al*, 2022). Thus, it is conceivable that the DNA-poor/HP1*β*-rich foci are PML bodies. In addition, we often noticed the association of a DNA-poor/HP1*β*-rich focus with a DNA-rich/HP1*β*-rich region (see e.g. Figure 1D1) and we wondered whether this represents examples of the early stages of formation of larger HP1-rich regions. PML bodies also often attach themselves to heterochromatin (Hayakawa *et al*, 2003), but we were unable to observe any complete merging of foci in live-cell time-lapse imaging of HP1*β* (see e.g. Video S1).

In addition to studying the mechanisms of HP1*β*-mediated heterochromatin formation we wanted to understand how HP1*β* may alter heterochromatin function. Surprisingly, we found that chromatin in DNA-rich/HP1*β*-rich regions is more dynamic than that within DNA-rich/HP1*β*-poor foci even though it is more densely packed in the former – in fact chromatin in DNA-rich/HP1*β*-rich foci exhibits similar dynamics to that in DNA-poor/HP1*β*-rich foci and euchromatin (Figure 6). By contrast, DNA is less dense, and diffusion of chromatin is significantly reduced in the DNA-rich/HP1*β*-poor foci (Figure 6A). Thus, it appears that the mechanisms of chromatin compaction in HP1*β*-rich vs HP1*β*-poor heterochromatin might be different.

There are two main models for repression of transcription by heterochromatin structure: proteins either compact nucleosomes into a folded structure that restricts access to the underlying DNA by the transcription machinery (the “packaging” model); or proteins cross-link DNA into a mesh-like structure that cannot be accessed by the transcription machinery (the “compartment” model) (Paro, 1993). The finding that DNA is very significantly more compacted in the DNA-rich/HP1*β*-rich foci, but that this does not restrict chromatin diffusion or prevent access to diffusing molecules (at least those of the size of HP1*β*), suggests that HP1*β* functions to compact chromatin fibres locally whilst leaving the space between the chromatin fibres accessible. This could be functionally important, e.g. to allow the reversible switching on/off of transcription of genes located in heterochromatin (Lu *et al*, 2000). On the other hand, the lower chromatin mobility and accessibility to diffusing proteins of DNA-rich/HP1*β*-poor foci might suggest that other proteins (e.g. histone H1) may instead function to cross-link chromatin fibres to form a mesh-like structure with a restricted pore size. This steric hindrance may provide a more repressive chromatin environment. The finding that histone H3K9me3 is depleted in DNA-rich/HP1*β*-poor foci compared to euchromatin (Figure 2E) is consistent with this idea, suggesting that the methylation machinery might have restricted access to these regions of densely packed chromatin.

It is thought that HP1*β* is essential for the maintenance of pluripotency in ESCs and for proper differentiation (Mattout *et al*., 2015), and we particularly wanted to understand how heterochromatin structure changes as mouse ES cells start to differentiate. To this end we studied an *in vitro* model where cells are maintained in naïve pluripotency by the presence of 2i and LIF. Upon removal of 2i and LIF, as mESCs exit from naïve pluripotency, we found that the level of HP1*β* increases concomitantly with a decrease in histone H3K9me3 binding sites. In 24 hr cells this results in a weaker enrichment (relative to euchromatin) of HP1*β* in DNA-rich/HP1*β*-rich heterochromatic foci (Figure 7D). Unexpectedly, by 48 hours after withdrawal of 2i/LIF, as cells enter a primed state, without any further changes in either global HP1*β* or histone H3K9me3 levels, HP1*β* rebinds and its localisation pattern becomes more like that of the DNA, both globally (Figure 7E) and at high resolution within DNA-rich/HP1*β*-rich heterochromatic foci (Figure 7G).

Thus, we propose the following model that is consistent with our observations (Figure 7H). The DNA-poor/HP1*β*-rich foci contain little chromatin, but many of the nucleosomes are modified with H3K9me3, which seeds the formation of an HP1*β* condensate (Figure 7H, left). Within the condensate HP1*β* exists in a range of states characterised by different degrees of mobility, being involved in a variety of stable and transient interactions with chromatin/other potential members of the condensate. This leads to a high fraction of apparently immobile chromatin-bound HP1*β* that is beyond what would be expected from a simple bimolecular reaction model, as well as to retention of apparently mobile HP1*β* molecules within this liquid phase. Overall, such a system relies on a “basal” layer of chromatin-bound HP1*β*, and thus many aspects of its behaviour are consistent with the stable chromatin binding model, but it also exhibits cooperativity and features of LLPS (Mittag & Pappu, 2022). In addition, as discussed above, the high local concentration of HP1 could lead to shadow chromodomain-mediated recruitment of Lysine-9 histone H3K9 methyltransferases, and the resulting positive feedback loop would also be expected to contribute to the apparent cooperativity.

The DNA-rich/HP1*β*-poor regions contain chromatin that is cross-linked and condensed in a HP1*β*-independent manner, e.g. through histone H1 binding and/or direct nucleosome-nucleosome contacts (Thoma & Koller, 1977), (Dorigo *et al*, 2003). These regions do not contain a high proportion of H3K9me3-modified nucleosomes, but due to chromatin condensation the local concentration of H3K9me3 is also somewhat elevated compared to the surrounding euchromatin. This too leads to a higher percentage of chromatin-bound HP1*β* molecules within this environment compared to euchromatin. However, due to crowding and the low mobility of the cross-linked chromatin, mobile HP1*β* (and potentially other proteins and complexes) cannot access these regions. Overall, the HP1*β* concentration does not reach the necessary threshold to nucleate a secondary condensate (Figure 7H, middle).

In naïve pluripotent ES cells major satellite transcription is high, which results in chromocentres having a different morphology and biophysical properties to more differentiated cells (Novo *et al*., 2022). On the other hand, super-resolution imaging studies have uncovered a segregation between DNA dense chromatin and RNA-rich regions as a major feature of genome organisation (Miron *et al*., 2020), (Hilbert *et al*, 2021). We found that the DNA-rich/HP1*β*-rich foci are heterogeneous in naïve mESCs, containing local areas that resemble either the DNA-poor/HP1*β*-rich or the DNA-rich/HP1*β*-poor regions (Figure 7H, right). We hypothesise that the segregation between these regions might be caused by the presence of large amounts of major satellite (or other) RNA in the DNA-poor/HP1*β*-rich pockets. In addition, active transcription within these foci might serve as a stirring factor leading to higher chromatin mobility (Gu *et al*., 2018).

In the first 24 hrs after exit from naïve pluripotency the increase in HP1*β* and the decrease in histone H3K9me3 levels, especially at major satellite and other repeats forming constitutive heterochromatin, leads to a loss in HP1*β* from heterochromatin (Figure 7I). Our previous studies suggested that a loss of PRC1 binding leads to an increase in transcriptional bursting and increased chromatin stirring in the formative state, and the increased inter-chromosomal intermingling and the generation of multiway chromatin hubs that we observed in the 3D genome structures (Lando *et al*., 2024).

However, the results presented here suggest that it is plausible that the loss in HP1*β* from heterochromatin might also contribute to these structural changes. We propose that after 48 hrs of differentiation, major satellite repeat (and/or other RNA) transcription might be downregulated, leading to a diminished RNA-rich heterochromatic phase. This would then lead to a rearrangement of heterochromatin structure into a more canonical form when HP1*β* is recruited back to chromatin. Studying the functional and mechanistic interplay between chromatin, transcription, RNA, and HP1 in this context will be an exciting avenue for future research.

## Acknowledgements

We thank Tessa Kretschmann for preparing the figures for publication, Luke Lavis (HHMI, Janelia Farm) for providing the Halo-tagged JF dyes, the Gurdon (Alex Sossick), CSCI (Peter Humphreys and Darran Clements) and Cambridge Advanced imaging Centre imaging facilities, and the CSCI DNA sequencing (Maike Paramor and Vicki Murray) and School of Biological Sciences flow cytometry (Joanna Cerveira) facilities. We thank the Laura Machesky group for use of their Airyscan microscope and the BBSRC (BB/R000395/1) for funding the STED microscope. We thank the Medical Research Council (MR/P019471/1 EDL) and the Wellcome Trust (206291/Z/17/Z EDL) for programme funding. We also thank the MRC (MR/R009759/1 BDH, and MR/M010082/1 EDL), for project grant funding, and we thank the Wellcome Trust/MRC for core funding (203151/Z/16/Z) to the Cambridge Stem Cell Institute. We are also grateful to the BBSRC and MRC DTP programs for funding AO’s and DS’s PhD studies.

## Author contributions

AO and EDL designed the experiments. DL and MW designed and constructed the mES cell lines where we fused a HaloTag to the endogenous *Cbx1* gene. NR, WB, XM and BDH carried out the ChIP-seq data collection, processing and analysis. AP, ZZ and DK designed, built and maintained the microscopes used for the live cell single molecule tracking and eSPIM experiments. WB developed the code to generate single molecule tracks from the localisation data. AO, DS, XM and ML carried out the STED imaging experiments. AO and DS carried out all the other imaging experiments and AO analysed the data with input from KU-J, BDH, DK and EDL. AO and EDL wrote the manuscript with assistance from all the other authors.

## Competing interests

The authors declare no competing interests.

## Materials and Methods

### Cell culture and line generation

Female 129/CAST F1 hybrid mouse ES cells (Joost Gribnau) were used throughout this study. Cells were routinely grown under serum-free 2i/LIF conditions (Ying *et al*, 2008) and typically passaged every second day. The media did not contain phenol red to decrease phototoxicity during live-cell imaging.

To create the HP1*β*-H(N) cell line, the endogenous *Cbx1* gene was genetically engineered using the CRISPR-Cas9 technology to produce an N-terminal fusion of HP1*β* with a HaloTag (Figure S1A). Genotyping PCR (using CACCTTGCCCTTGACAACTCG as the forward and TATCCAGCCAGGAAATGTGCC as the reverse primer) and western blot analysis indicated that the resultant cell line was a mixture between homozygotes and hemizygotes, with the latter having one of the *Cbx1* alleles successfully targeted and the other knocked out (Figure S1B,C). While the homozygotes expressed the same level of HP1*β* as wild-type cells, expression was two times lower in the hemizygotes (Figure S1D). HP1*β* in these cells was labelled with JF646, and the genotypes were separated by double fluorescence-activated cell sorting (FACS), using side scatter to account for the cell cycle effects (Figure S1C,D). Protein level measurements using flow cytometry and genotyping PCR were used to confirm the purity of the resultant populations.

### RT-qPCR

Reverse transcription-quantitative PCR (RT-qPCR) experiments were performed to assess the RNA levels of genes known to be deregulated upon *Cbx1* knockout (Mattout *et al*., 2015) and common pluripotency markers. RNA was extracted using the standard TRI reagent protocol (Sigma T9424) and a two-step RT reaction was performed alongside no polymerase controls. QPCR was carried out in triplicate with either TaqMan probes (Life Technologies) or SYBR Green mix (Life Technologies 4385614) and custom-made primers (Table 1), and for each gene a no-template control was also included. The levels of the test gene mRNA and the reference gene mRNA (*Gapdh*, *Ppia*, *Atp5a1* – all gave similar results) in wild-type and targeted cells (three biological repeats) which were compared using double delta Ct analysis and statistical testing.

**Table 1:**
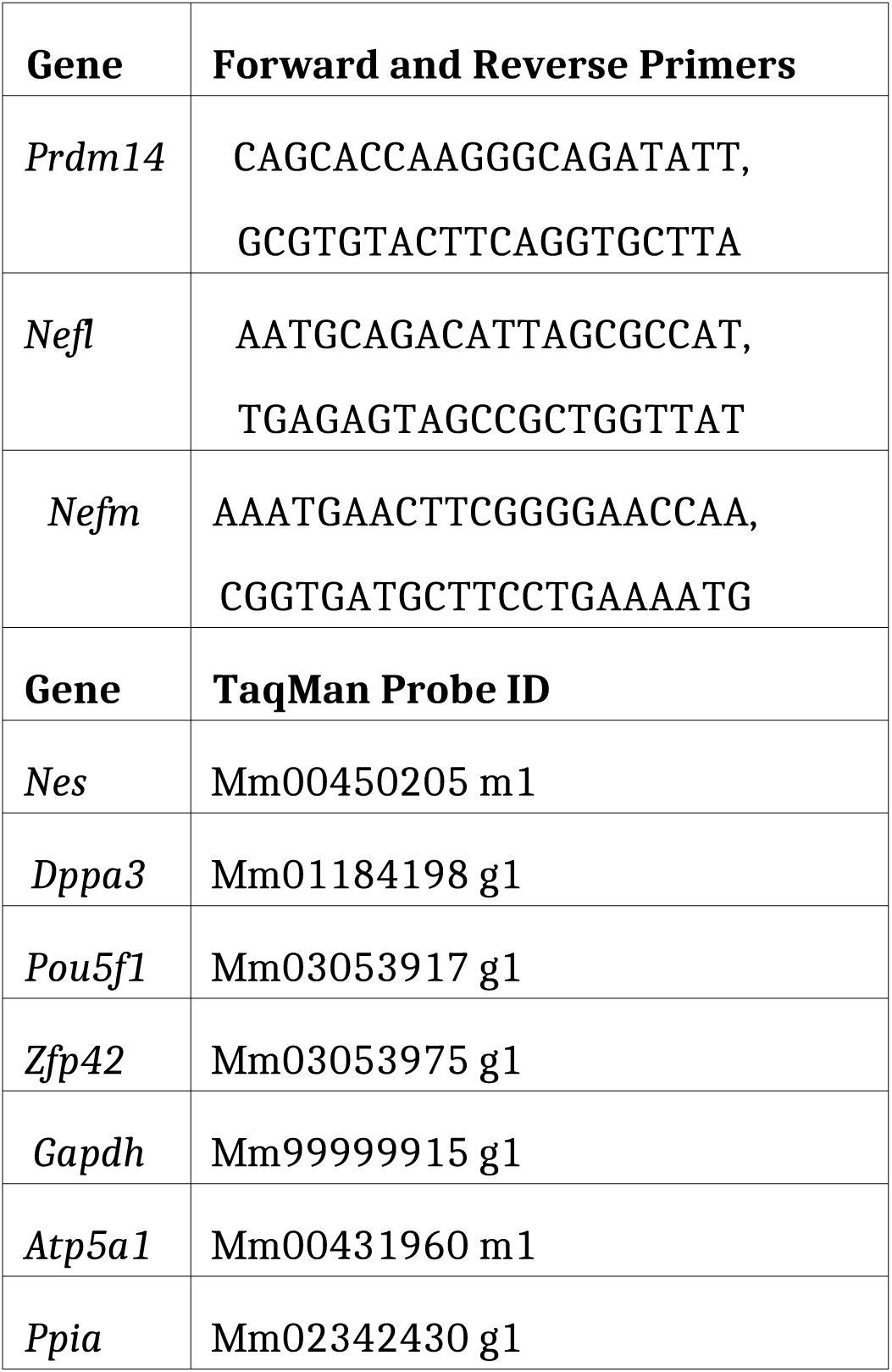
RT-qPCR primers and probes.

### Western blot

Nuclear protein extraction and western blotting were carried out in a standard manner, using the anti-HP1*β* mouse mAb (Invitrogen MA3-053, 1:2000) and anti-RNAPII NTD rabbit mAb (Cell Signalling 14958S, 1:2000; loading control) as primary antibodies, and sheep anti-Mouse IgG + HRP (GE Healthcare NA931V, 1:2000) and donkey anti-Rabbit IgG + HRP (GE Healthcare NA934V, 1:2000) as secondary antibodies. The bands were visualised on film with an HRP detection kit (Fisher Scientific 10340125).

### Alkaline phosphatase assay

For this experiment, mESCs were transferred into serum/LIF medium 7-9 days before scoring. Two days prior to this, ∼30,000 cells were plated onto wells of a 6-well plate. Cells were fixed according to standard protocols and stained for alkaline phosphatase activity using the Sigma 86R-1KT kit (without counter labelling with haematoxylin). Approximately 100 colonies were scored using a bright-field light microscope, according to their labelling (clear pink labelling, no labelling, or mixed/unclear). Scoring was performed blind. The experiment was conducted twice for each cell line.

### Sample preparation for fixed-cell microscopy

MESCs were grown on glass-bottom 35 mm dishes (MatTek #1 or #1.5), coated with poly-L-ornithine and laminin. To improve dye and antibody penetration, ESCs were seeded sparsely and grown for only one day, giving rise to small-sized colonies. Cells were fixed with 4% formaldehyde for 10 min at room temperature, gently permeabilised with 0.1% Triton-X100 in PBS for 4 min (JF dyes) or 10 min (AF594) and blocked with 1% BSA in PBS for 20-60 min to improve dye penetration and specificity (Grimm *et al*, 2017). Samples were stored and imaged in PBS.

HP1*β*-HaloTag was labelled with HaloTag ligand-conjugated Janelia Fluor 549 (H-JF549; gift from Luke Lavis), Janelia Fluor 646 (H-JF646; gift from Luke Lavis) or Alexa Fluor 594 (H-AF594 for STED). The latter was produced from AF594 NHS ester (Invitrogen) and HaloTag ligand (Promega) which were conjugated and purified by HPLC in-house. JF dye labelling was performed in live cells by incubation with 250 nM of the dye for 15 min at 37 C, followed by three 15 min media washes with quick PBS rinses in between. AF594 labelling was performed after fixation and permeabilisation using a 1 hr incubation with 250 nM of the dye, followed by 2 washes in PBS.

To visualise DNA, 1 g/ml Hoechst33342, 500 nM SPY505-DNA (Spirochrome), 500 nM SiR-DNA (Spirochrome) or 50 nM JF464-Hoechst (gift from L. Lavis; for STED) were added after other labelling procedures to the sample at least 1 h before imaging.

### Immunofluorescence (IF)

After fixation, cells were permeabilised (1% TritonX-100, 10 min), blocked (10% BSA, 1h), incubated with primary antibody (in 10% BSA, 4-5 hrs at room temperature or overnight at 4 C) and washed 3 times (0.1% TritonX-100, 0.2% BSA, 10 min each time). Next, the sample was labelled with the secondary antibody (in 10% BSA, 1hr), washed again three times, re-fixed (3% formaldehyde, 10 min) and rinsed with PBS and water. When combined, JF dye labelling was performed before IF, while AF594 and DNA labelling was performed after IF, with the gentle permeabilisation replaced by the harsher IF protocol.

The antibodies used for IF are listed in Table 2.

**Table 2:** List of antibodies used for immunofluorescence.

| Antibody | Supplier and cat. no. | Dilution |
| --- | --- | --- |
| **Primary antibodies** |  |  |
| Anti-HP1 $\beta$ Rb pAb | Abcam ab10478 | 1:200 |
| Anti-HP1 $\alpha$ Rb mAb | Abcam ab109028 | 1:250 |
| Anti-H3K9me3 Rb pAb | Active Motif 39062 | 1:500 |
| **Secondary antibodies** |  |  |
| Dk anti-Rabbit IgG + Cy3 | Jackson ImmunoResearch Conjug in-house to Cy3 (GE Healthcare) | 1:150 |
| Dk anti-Rabbit IgG + AF594 | Invitrogen R37119 | 1:400 |

### Airyscan confocal, laser scanning confocal and STED microscopy

The Airyscan confocal imaging was performed on Zeiss LSM900 Airyscan2 microscope at Cambridge Advanced Imaging Centre or a Zeiss LSM980 Airyscan2 microscope in the University of Cambridge, Department of Biochemistry, both equipped with 405 nm, 488 nm, 561 nm and 640 nm lasers. The pixel size and z stack steps optimal for highest resolution were selected automatically. The scan speed was set to maximum (6-9, ∼1 s pixel dwell time), laser intensities were 0.5-2%, and 2-4 line averaging was chosen. Different channels were acquired sequentially frame-by-frame to minimise crosstalk. Images were processed with 3D Airyscan processing and weak deconvolution within the Zen Blue software.

Laser scanning confocal imaging was also performed on the Zeiss LSM980 Airyscan2 microscope in the University of Cambridge, Department of Biochemistry, with scan speed 14 and other settings similar to above. Weak deconvolution was also applied to these images, using the Zen Blue software.

Two-colour 3D STED microscopy was conducted on an Abberior 3D STED microscope at the University of Cambridge, Department of Genetics. The 561 nm (for AF594) and 640 nm (for JF646) excitation, as well as the 775 nm depletion lasers were aligned prior to every imaging session. Furthermore, a high degree of correspondence between channels was ensured by the employment of the same depletion laser. The channels were acquired sequentially line-by-line, and the absence of significant crosstalk was confirmed using appropriate controls. 20-by-20 nm pixel size was chosen for maximum resolution. The laser intensities and pixel integration time were optimised empirically, with the typical values being 50% (561 nm), 20% (640 nm) and 30% (775 nm), and 20-40 μs pixel integration time with a line averaging of 5-8. A one pixel-wide Gaussian filter was applied to the images.

### Airyscan confocal and laser scanning confocal image analysis

Image analysis was performed in Fiji. The results shown in Figures 1, 5 and 7 were obtained using maximum projections of z-stacks; and those shown in Figure 2 were obtained from single slices.

**1. Segmentation of nuclei and colocalisation analysis.** First, the individual nuclei were segmented by applying the Li global thresholding algorithm to the sum of the normalised HP1*β* and DNA images. The segmentation was manually checked and corrected when necessary. The colocalisation analysis was performed by calculating the 2D Pearson’s correlation coefficient between different channels on a per-nucleus basis, using the Coloc 2 plug-in.
**2. Segmentation of foci** The DNA and HP1_β_ foci were then segmented independently in the respective channels, using the Max Entropy and Triangle thresholding algorithms, respectively (Figure S3A). The former was performed in each nucleus separately, while the latter was applied to all nuclei in the field of view at the same time. Segmented particles that were too small were discarded. The resultant masks were also checked and corrected manually. In particular, small near-round bright HP1*β* puncta and adjacent larger, less bright (and usually heterochromatic) foci were separated.
**3. Classification of foci.** Since the shape of the foci in the two channels can be different, and sometimes foci can be touching but distinct, it was hard to design an automated procedure for annotating the three types of foci. Thus, the DNA-dense foci were marked as either HP1*β*-enriched or non-enriched, and the HP1*β*-dense puncta were annotated as either DNA-rich or DNA-poor manually. If the foci overlapped at least partially, they were considered DNA-rich and HP1*β*-rich (“canonical heterochromatin”). However, small overlaps of the segmented regions were ignored when it was evident from the raw image that the foci were close but non-overlapping.
**4. Quantification** It was possible to count the euchromatic HP1*β* puncta per cell, since they were small and distinct. However, counting would be meaningless for more extensive, heterogeneous and connected/overlapping DNA-dense regions. Hence, e.g. Figure 1B shows the area of HP1*β*-rich heterochromatin divided by the total area of heterochromatin per cell. The segmentation was then used to measure the mean intensity of HP1*β*, DNA and H3K9me3 signal within each focus and in euchromatin (defined as the area outside both HP1*β* and DNA condensates). The values for euchromatin were used for normalisation. The measurements were analysed and plotted in R.

### STED signal correlation

Correlation between channels within foci in STED images was calculated in Fiji. To this end, foci were first segmented by applying Max Entropy thresholding to a sum of the normalised channels (Figure S3B). Foci that were too small were discarded, and the resultant mask was manually checked. The segmented regions were then dilated by 5 pixels to include the border of the focus. 2D Pearson’s correlation coefficient was computed per focus, using the Coloc 2 plugin, and the results were then analysed and plotted in R.

### Statistical analysis of microscopy data

The statistical analysis for Airyscan, confocal or STED microscopy-derived measurements, such as inter-channel Pearson’s correlation coefficients (Figures 2B, 7B, 7G), absolute or relative intensities of individual foci (Figures 2D, 2E, 5B, 7D), intensity of euchromatin per cell (Figure 7D) or number of foci per cell (Figure 5C), was performed as follows. First, the distributions of the relevant metric from 2-3 individual biological repeats (except for the STED experiments, which were primarily used to confirm the conclusions from the other imaging studies) were compared to confirm the reproducibility of the measurements. In Figure 2B, there were substantial differences in the experimental techniques used to perform the different replicates, and thus each replicate was analysed separately, yielding similar conclusions. In Figure 5B, the absolute fluorescence intensities in the two replicates differed due to different efficiency of staining, but this effect was corrected for by normalisation to the intra-replicate mean. In all other cases, the results were reproducible, and thus the data from the different biological replicates was pooled.

Statistical comparisons were made between the pooled distributions to test whether the results were statistically significant given the inherent biological variation. The normality of the distributions was examined using Q-Q plots and the Shapiro-Wilk test, and the equality of variance between the samples was tested using Bartlett’s or Levene’s tests. The appropriate statistical test for comparison of means between samples was then selected based on the normality, sample size and equality of variance. The precise tests used for statistical analysis, as well as the sample sizes, are described in each figure legend.

### Sample preparation for SPT

For live-cell SPT, mESCs were seeded onto #1 glass-bottom dishes as described above. Prior to imaging, HP1*β* was labelled with 10 nm HaloTag ligand-conjugated spontaneously blinking dye Janelia Fluor 639b (H-JF639b, a gift from Luke Lavis (Holland *et al*, 2024)) in a 15 min incubation at 37 C. After three 15 min washes with media, 500 nM SPY505-DNA was added to the media and incubated for at least 1 hr prior to imaging.

### SPT image acquisition in live cells

A custom-built inverted fluorescence microscope, based on the Nikon Eclipse Ti2 body with a Perfect Focus system, was used in this study. The stage and part of the microscope body were encased in an incubator to maintain live cells at 37 C. The 488 nm and 638 nm laser beams (200 mW 06-MLD, Cobolt; 360 mW L2C laser combiner, Oxxius) with excitation filters (68840, Edmund Optics; FF01-637/7-25, Semrock) passed through the adjustable neutral density filters, and were then expanded and collimated with Galilean beam expanders, and combined using dichroic mirrors. A 60x oil immersion objective (CFI Apochromat TIRF 1.49 NA, Nikon) was used to focus the illumination onto the sample and to collect the emitted light. The laser intensities entering the sample were measured to be approximately 10 kW/cm^2^ (488 nm) and 40 kW/cm^2^ (638 nm), when no neutral density filters were used. Highly inclined and laminated optical sheet (HILO) illumination was employed to decrease out-of-focus background, and further to this end the diameter of the 638 nm beam was minimised (∼20 μm). An UltraFlat quad-band dichroic mirror (ZT405/488/561/640rpc-UF3, Chroma) was used to separate excitation and emission, and a Köhler lens focused the light onto the back focal plane of the objective. Within the microscope body, the light passed though another built-in 1.5x lens. The emitted light was separated into two pathways using another dichroic mirror (FF624-Di01, Semrock) and, after passing through emission filters (67030, Edmund Optics; 67038, Edmund Optics + BLP01-635R-25), it was focussed onto EMCCD cameras (Evolve 512 Delta, Photometrics; EM gain 250) using tube lenses. The final pixel size was 140 nm.

The laser beams were aligned prior to every experiment, using a series of pinholes and a fluorescent bead sample. However, since the optical setup is not perfect, the beams that overlapped perfectly in the epifluorescence regime sometimes diverged slightly in HILO. Thus, both beads and double-labelled cells were used to confirm that the lasers illuminated the same plane. The cameras were also aligned in x, y and z using fluorescent beads, and two-colour images of beads were acquired to computationally correct for any remaining mismatch.

The red (SPT of HP1*β*+H-JF639b (Holland *et al*., 2024)) and green (wide-field imaging of DNA+SPY505) channels were imaged simultaneously. Continuous acquisition with a 3 ms exposure time, which was the maximum speed the cameras could achieve (with pre-sequence camera clearing), was performed to enable observation of fast-diffusing molecules. Since chromatin domains in ESCs move by a noticeable amount in a few minutes, each cell was imaged for only 2.5 min (40,000 frames). The SPT imaging had to fulfill the following criteria: a) to obtain a sufficient number and diversity of trajectories to investigate diffusion in different environments, and b) to collect enough localisations to reconstruct the HP1*β* distribution at high resolution. Super-resolution image reconstruction requires one to image as many distinct molecules as possible, with average densities of ∼0.5 localisations/μm^2^ per frame being most appropriate, best achieved with dense labelling and high laser intensities. On the other hand, accurate tracking requires localisations to be ∼5-10-fold sparser and to not bleach within a frame. Notably, in our case we did not want the trajectories to be too long, so that we could probe the behaviour of different populations of molecules in the same focus. Thus, the labelling density and the 638 nm laser intensity (∼6.5 kW/cm^2^) were optimised empirically, bearing these criteria in mind (Figure S4A). The initial imaging period with high numbers of localisations was used only for image reconstruction and not for diffusion analysis (see below). The 488 nm laser intensity was chosen as ∼0.01 kW/cm^2^, such that binning frames every 15 s resulted in a reasonably bright image (Figure S4A).

Two biological replicates were acquired independently using the same setup, with 15 and 21 cells, respectively – no cell death was apparent neither during imaging nor ∼5 min after.

### SPT controls

Immobilised H-JF639b dye was used as a control to investigate the limits of the technique. The latter sample was prepared by incubating a glass-bottom dish coated with poly-L-lysine with 50 nM of H-JF639b in PBS for 5 min, followed by a PBS wash. The sample, which had similar density of localisations to the live datasets, was then imaged in warm cell culture medium under the same conditions as live-cell SPT.

### SPT image and data processing

The localisations in the HP1*β* SPT channel were extracted using the GDSC PeakFit Fiji plugin (Etheridge *et al*, 2022). For image reconstruction, the first 100-1500 frames were excluded, because background fluorescence was stronger at the beginning of each sequence. To exclude multiple occurrences of the same molecule, localisations within a 300 nm radius in successive image frames were linked together in “tracks” using a custom Python programme (available at https://github.com/TheLaueLab/trajectory-analysis). The tracks were allowed to “skip” one frame to allow for the blinking behaviour of fluorophores thereby avoiding the artificial splitting of trajectories, which would lead to an artifactual increase in local density. The average xy-positions of each “track” then formed the image, and these points will further be called “SMLM localisations” to distinguish them from the SPT trajectories (Figure S4A,B).

For SPT, more frames at the beginning of the acquisition had to be discarded to minimise erroneous trajectory joining. For each cell, a threshold was selected, such that a) after it, localisations within the same frame closer than 400 nm would only occur <0.01 times per frame, b) the overall localisation density was <10 per frame, and c) the jump distance distributions from 10,000 frame-long batches were reproducible (Figure S4A). Typically, the first ∼10-15,000 frames were excluded. The localisations were then joined up into trajectories, using a 400 nm radius (larger radius possible due to sparsity) and not allowing any gaps, since splitting up a trajectory was less deleterious than joining two localisations erroneously. 50-200,000 SMLM localisations and 20-130,000 SPT trajectories per cell were obtained, with a total of 750,000 and 1,750,000 SPT trajectories for each biological replicate.

Every block of 5,000 image frames in the DNA image stacks were binned, and the linear transformation calculated from the fluorescent bead control was applied to ensure that the red and green channels corresponded to each other. A visual check revealed that the images were shifted by 1 pixel in x and y, which was not the case for the bead controls. This appeared to be a software bug and was corrected for.

### Foci segmentation and SPT trajectory classification

The DNA foci were segmented in Fiji. First, the stacks were upsampled 10 times in x and y and then the Top Hat morphological filter was applied. Small particles were removed from the resultant mask, and it was checked and corrected manually (Figure S4C). Each focus was then manually displaced into a clearly euchromatic (not nucleolar and not heterochromatic) area within the same cell, producing its corresponding “pseudo foci” mask.

The HP1*β* super-resolution image (SMLM localisations) was segmented using DBSCAN clustering in R (Figure S4B). A 50 nm radius was chosen; the minimum number of points within a cluster was set to be inversely proportional to the mean distance to nearest neighbour squared, with the optimal coefficient (6650) estimated empirically by testing several cells. “Pseudo clusters” were generated automatically by randomly shifting the cluster cores into non-clustered regions, retaining their size and shape (Figure S4B). Since the density of SPT data outside of clusters is naturally lower and the clusters are small, the “pseudo focus” sampling procedure was repeated 6 times and the results were pooled to generate sufficient data to make comparisons. The SPT points were counted as belonging to an HP1*β* focus/pseudo focus if they were not further than 50 nm from any SMLM point making up the DBSCAN cluster/pseudo cluster cores.

Because the HP1*β* puncta were much smaller and better-resolved than heterochromatic areas, they were classified as either DNA-rich or DNA-poor in a binary way. The HP1*β* foci was said to be DNA-rich, if >30% of SPT localisations within it also belonged to a DNA focus (Figure S4D). In other words, all the localisations within this HP1*β* cluster were marked as residing within a HP1*β*-rich and DNA-rich area. The rest of heterochromatic localisations were annotated HP1*β*-poor heterochromatin, while the rest of the HP1*β* foci were defined as euchromatic. Notably, the corresponding HP1*β* pseudo foci were then also annotated as “DNA-rich” or “DNA-poor” pseudo foci, allowing the geometry in the pseudo focus control to remain the same as in the real data.

Plotting the jump distance distributions revealed that when either only one end of the jump or both ends are required to be situated within a focus, the jump distance distribution of data from pseudo foci differed significantly from euchromatin overall (skewed to the right in the former case and to the left in the latter). A much smaller bias was observed when the coordinate of the midpoint of each jump was used for classification, and thus this strategy was employed (Figure S4D). Every step within a trajectory was thus classified as belonging to a compartment, pseudo compartment or euchromatin, as described above. If a trajectory transitioned between compartments, it was split, with the breakpoint duplicated.

### Diffusion analysis with SA-SPT

The SA-SPT algorithm (Heckert *et al*., 2022) was employed to investigate the full range of diffusive behaviours without prior assumptions about the underlying populations and accounting for localisation precision. For Figure 4A, 100,000 non-classified trajectories from live cell and control data, processed in the same way, were used. The localisation precision was generally found to be ∼20 nm. In subsequent panels, each class of trajectories was subsampled with replacement 100 times to equalise the sample sizes and to test the robustness of the result. Only the regular Brownian motion (with localisation error) likelihood function could be used, since the (classified) sample size was not large enough for the much larger parameter space of the fractional Brownian motion model.

For Figure 4D, the HP1*β* density within all foci and pseudo foci was defined as the 1/(4*NNdist2), where NNdist is the average nearest-neighbour distance between SMLM localisations within DBSCAN cluster cores or <50 nm from them (for both DNA-poor and DNA-rich HP1*β* puncta) or within the segmented DNA-rich foci. The values averaged across all foci/pseudo foci were then used for the calculation.

### Spinning-disk confocal live-cell imaging

Mouse ESCs for this experiment were seeded as above, and HP1*β* and DNA were labelled with 250 nM JF549 and 500 nM SiR-DNA, respectively, as described above. The data for Hi-D chromatin diffusion analysis was acquired on a Nikon CSU-W1 SoRa spinning disk confocal microscope at the Gurdon Institute Imaging Facility. The 60x oil objective with 4x extra magnification and SoRa boost were used to obtain maximum resolution. Imaging was performed sequentially in the yellow and red channels, with the 561 nm (20%) and 638 nm (20%) lasers used for illumination. The exposure time was set to 200 ms, so that the effective time resolution (2.5 Hz) was similar to the original study (Shaban & Seeber, 2020), and 150 frames were recorded per field of view (1 min total acquisition time). During acquisition, the fluorophores bleached by <5%. The live cells were kept at 37 degrees C during imaging.

### Processing of live-cell imaging with Hi-D

The DNA data was analysed using the published Hi-D software(Shaban & Seeber, 2020) . First, the nuclei were segmented, and the nucleolar regions were excluded from analysis. Next, for each pixel, the optical flow-based Hi-D method was used to fit the five diffusion models (regular Brownian or anomalous diffusion, with or without a directional component), select the best-fitting one and extract the parameters. The output was then further analysed in R. The apparent diffusion coefficient values for the pixels where the anomalous model was selected were much higher than for those where the simple diffusion model was preferred. Thus, it appeared that in the latter case, confinement influenced the estimated diffusion coefficient strongly. This rendered these two types of pixels non-comparable and, given that chromatin is expected to display confined diffusion, only pixels with the anomalous diffusion model were kept for further analysis. The directional component, on the other hand, had a minimal influence on the diffusion coefficient. It was visually confirmed to correspond to coordinated chromatin domain motion and was not considered further.

The values of the chromatin diffusion coefficient per pixel were then correlated with the brightness of the pixel in the DNA and HP1*β* channels. Heterochromatin and euchromatin were defined as pixels with DNA signal >1.5 or <1.1 the nuclear average, while HP1*β*-enriched and -depleted regions were defined as being >1.4 or <1.1 brighter than the mean, respectively.

A fixed-cell control was prepared, and the data was acquired and processed analogously. In addition, live data with varying laser intensities was acquired. Both controls indicated no effect of the brightness on the diffusion coefficient estimates within the range tested, suggesting that the differences in signal intensity between foci and euchromatin did not affect the results.

### eSPIM light-sheet live-cell imaging

The live mouse ESCs for this imaging were plated and labelled with 250 nM JF549 (HP1*β*) and 500 nM SiR-DNA (DNA), as described above.

An open-top single-objective light sheet microscope based on the Oblique Plane Microscopy (OPM) configuration was employed for the experiments. The overall design followed a similar layout to that described by (Yang *et al*, 2019). A primary 60× NA 1.27 water-immersion objective (CFI Plan Apo IR 60XC WI, Nikon), mounted on an inverted microscope frame (Eclipse Ti-U, Nikon), was utilised for both light sheet excitation and fluorescence collection. Two continuous-wave diode lasers, 561 nm (100 mW, 06-MLD-561, Cobolt) and 638 nm (180 mW, 06-MLD-638, Cobolt), were expanded, collimated, and combined into a single optical path. After combination, the beam was shaped into a 1D Gaussian profile by passing through a cylindrical lens. A quad-band dichroic mirror (Di01-R406/488/561/635, Semrock) was used to separate excitation and emission light. The excitation beam was directed through two 4f optical relays before being focused at the primary objective’s back focal plane (BFP), forming an approximately 1.8μm-thick light sheet tilted at approximately 30 degrees. Volumetric imaging was achieved by rapidly scanning the light sheet using a 1D galvanometric mirror (GVS201, Thorlabs), which was driven by a custom LabView script.

Fluorescence emission collected by the primary objective was transmitted through the same dichroic mirror and relayed onto a 40× air objective (CFI Plan Apo Lambda D Air, NA 0.95, Nikon), with the pupil plane matched to that of the primary objective. The emission was then captured by a custom-designed glass-tipped tertiary objective (NA 1, AMS-AGY v1.0, Calico Lab) positioned at a 30-degree angle relative to the optical axis. A 321 mm tube lens was used to project the image onto a sCMOS camera (Prime 95B, Photometrics). A three-channel optical splitter (OptoSplit III, Prior) was integrated into the system to enable simultaneous multi-colour imaging. During imaging, the sample was kept at 37°C in 5% CO_2_.

System control, including camera acquisition, shutter operation, and sample stage movement, was performed using the open-source software Micro-Manager 2.0. Image stacks were acquired during light sheet scanning across the sample, with a 526 nm step size between the tilted slices. 10 different positions were imaged for >3.5 hrs each, with a 70s interval. For multi-colour alignment, calibration images of 100 nm TetraSpeck™ fluorescent microspheres (T7279, ThermoFisher) adhered to the coverslip were used to quantify and correct translational and rotational offsets between channels. Deskewing and 3D reconstruction of the raw data were carried out using a custom Python pipeline available at https://github.com/Zui409/3D-Deskew-and-Reconstruction-for-single-objective-SPIM-data, resulting in a voxel size of 120x120x120 nm^3^. Lucy-Richardson deconvolution (10 iterations) was performed on the processed images. One of the resultant stacks was used to create Video S1 in Fiji.

### Quantification of H3K9me3 histone modification abundance at satellite and other types of repeats

The mouse mm10 repeats annotation was downloaded from UCSC. Mouse ES cell H3K9me3 ChIP-seq data at early differentiation time points (from GEO GSE214264, (Lando *et al*., 2024)) was re-processed following the RepEnrich guidelines (https://github.com/nskvir/RepEnrich). More specifically, data was mapped to the mm10 genome using bowtie, allowing multi-mapping. Ambiguously mapped reads were output to fastq files using the --max option, while uniquely mapped reads were kept in separate files. The RepEnrich python script was run on the re-processed data with the --pairedend TRUE option. H3K9me3 abundance at major/minor satellite and other different types of repeats was then retrieved from RepEnrich output.

## Notes

### Competing Interest Statement

The authors have declared no competing interest.

