## Supplementary document S1 - Supplementary Figures S1-S7 for "HP1*β* and densely packed chromatin form separate microdomains in mouse ES cells, which are reconfigured upon exit from naïve pluripotency"

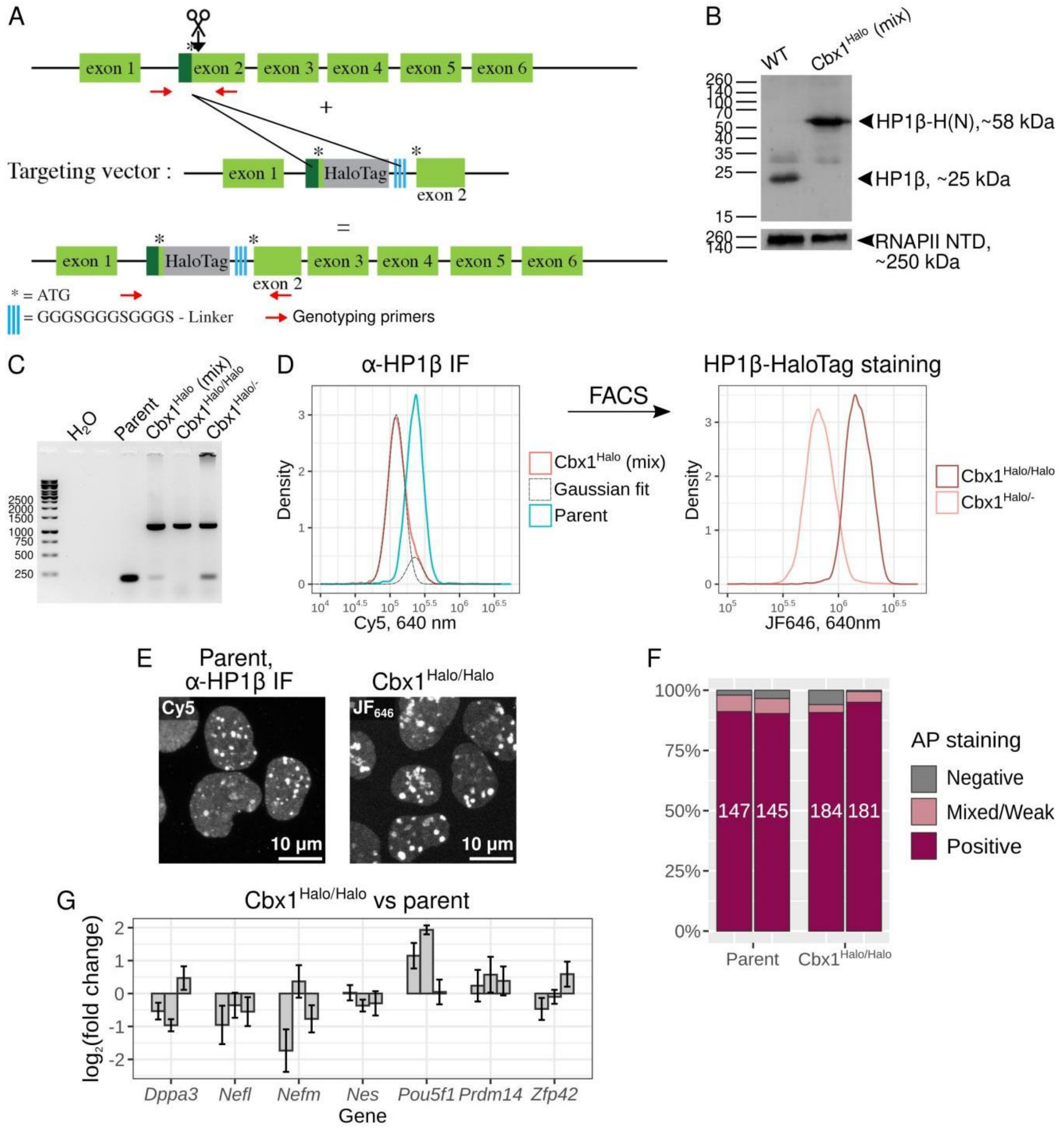


**Figure S1. Construction and characterization of *Cbx1$^{Halo/Halo}$* and *Cbx1$^{Halo/-}$* mESC lines, related to STAR Methods.** A. Targeting the *Cbx1* gene with CRISPR-Cas9. The repair template generated cells encoding a N-terminal HaloTag-$(GGGS)_3$ linker-HP1β fusion protein. B. Western blot for HP1β indicating that no wild-type protein is present in the knock-in cell line, which turned out to be a mixture of homozygous and hemizygous targeted cells. C. Genotyping PCR showed a faint wild-type band in the original targeted cell line (wild-type allele band at 200 bp and targeted allele band at 1130 bp). D. Immunofluorescence against HP1β combined with flow cytometry (left) indicated that the cell line was a mixture of cells expressing HP1β at endogenous and at two-fold lower levels. These cells were separated by FACS, and JF646 labelling and flow cytometry (right) confirmed that the two populations stably express different amounts of HP1β. Genotyping PCR (C) showed that the high-expressing population was homozygous at the targeted locus, while the low-expressing population had an untargeted allele that did not produce stable protein. E. The localisation of HaloTag-HP1β in the nucleus was like that in the parent cell line (visualised by immunofluorescence). Shown are maximum projections of confocal z stacks, where the intensity is adjusted for clarity. F. Alkaline phosphatase assay of the wild type and HP1β-Halo(N) cells, grown in serum/LIF conditions for three passages. Each bar corresponds to a biological replicate (two per cell line), with the number of colonies counted shown in the middle. The lack of any difference between the cell lines suggests that HP1β function is not compromised by the HaloTag (Mattout *et al.*, 2015). G. RT-qPCR analysis of RNA levels of genes that are misregulated upon HP1β knockout (Mattout *et al.*, 2015), together with the pluripotency markers *Pou5f1* and *Zfp42,* in *Cbx1*$^{Halo/Halo}$ cells compared to the parent cell line. *Gapdh* was used as a reference gene. Each bar shows a biological replicate, with standard deviation of technical replicates shown as error bars. None of the genes displayed significantly different RNA levels (two-tailed t-test).

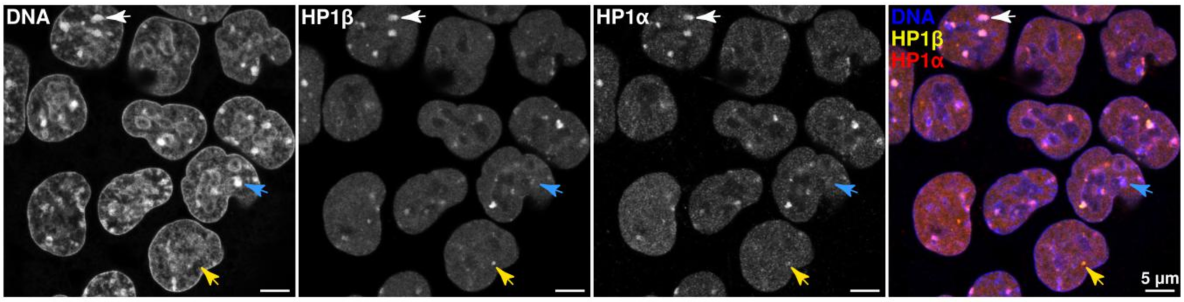


**Figure S2. Colocalisation of HP1β and HP1α, related to Figure 1.** Airyscan confocal images of DNA (Hoechst), HP1β (JF646-Halo) and HP1α (IF, Cy3) in mESCs (single slice). Examples of DNA-poor/HP1β-rich/HP1α-rich (yellow arrow), DNA-rich/HP1β-rich/HP1α-poor (blue arrow) and canonical heterochromatic DNA-rich/HP1β-rich/HP1α-rich foci (white arrow) are indicated.

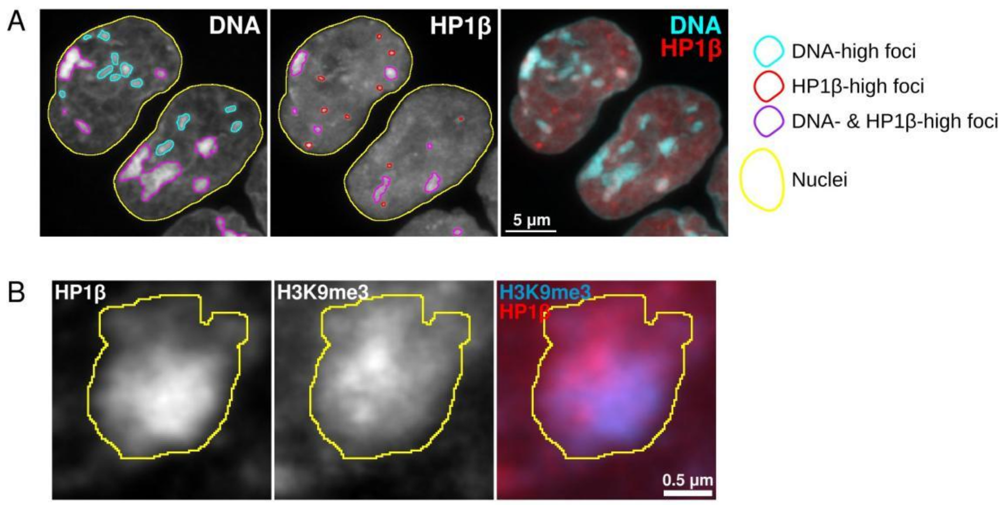


**Figure S3. Examples of segmented nuclei and foci from microscopy images, related to Figures 1, 2 and 3, and STAR Methods.** Examples of segmentation of nuclei and either DNA or HP1β foci from Airyscan confocal images (maximum projection) (A), or of a large HP1β-rich and H3K9me3-rich focus from a STED image (single slice) (B).

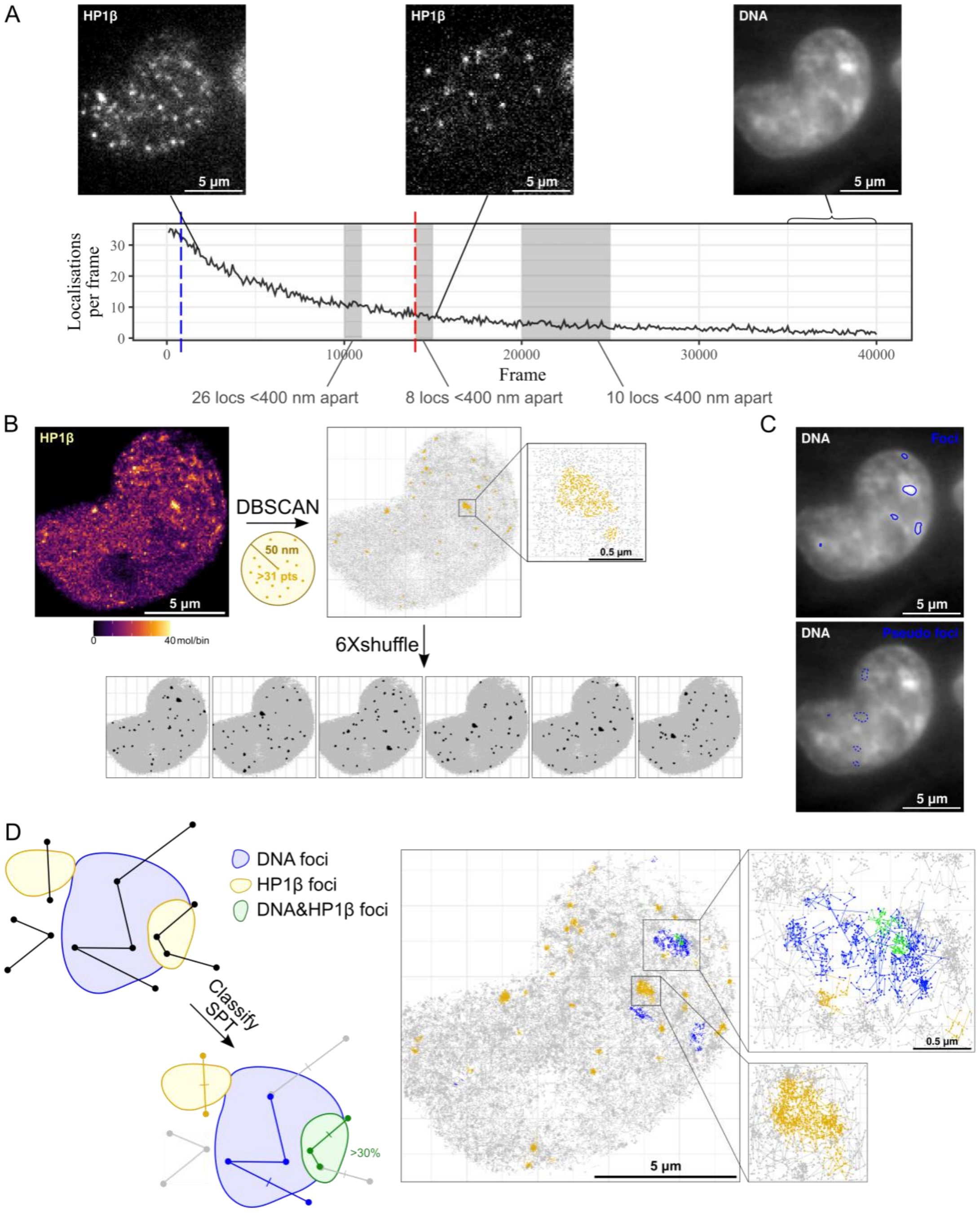


**Figure S4. Processing of SPT data, related to Figure 4 and STAR Methods.** A. Example of SPT data and its processing. Based on the number of localisations per frame (graph) and the number of localisations that are within the same frame and closer together than 400 nm – the number of such localisations within the grey-shaded periods is shown – the frames after frame 14,000 were used for SPT diffusion analysis (red dashed line). For super-resolution image reconstruction, all images after frame 800 were used (blue dashed line). Representative frames, as well as a binned DNA image, are shown. B. The super-resolution image of HP1β was segmented using DBSCAN (50 nm radius, minimum 32 points for this cell), and the resultant clusters were then shuffled to create pseudo foci. C. Example of segmented DNA image (top) and the corresponding shuffled pseudo foci (bottom). D. Left, each jump in a trajectory was classified based on its midpoint location within or outside the segmented foci (thin short lines shown for some jumps for illustration). The HP1β foci that had a >30% overlap with the DNA foci were defined as DNA-rich/HP1β-rich. Right, an example segmented SPT data.

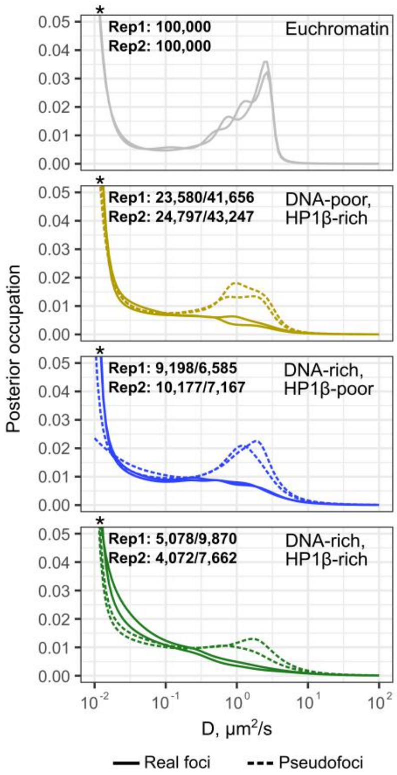


**Figure S5. Spectra of apparent diffusion coefficients for HP1β in euchromatin and different nuclear foci and their corresponding control pseudo foci, related to Figure 4.** The distributions were produced analogously to those in Figure 4B, but without subsampling. The number of trajectories in each environment in each of the two biological replicates is shown in the top-left corner of each panel (real foci/pseudo foci). The *'s show that the spectrum extends beyond the plot area.

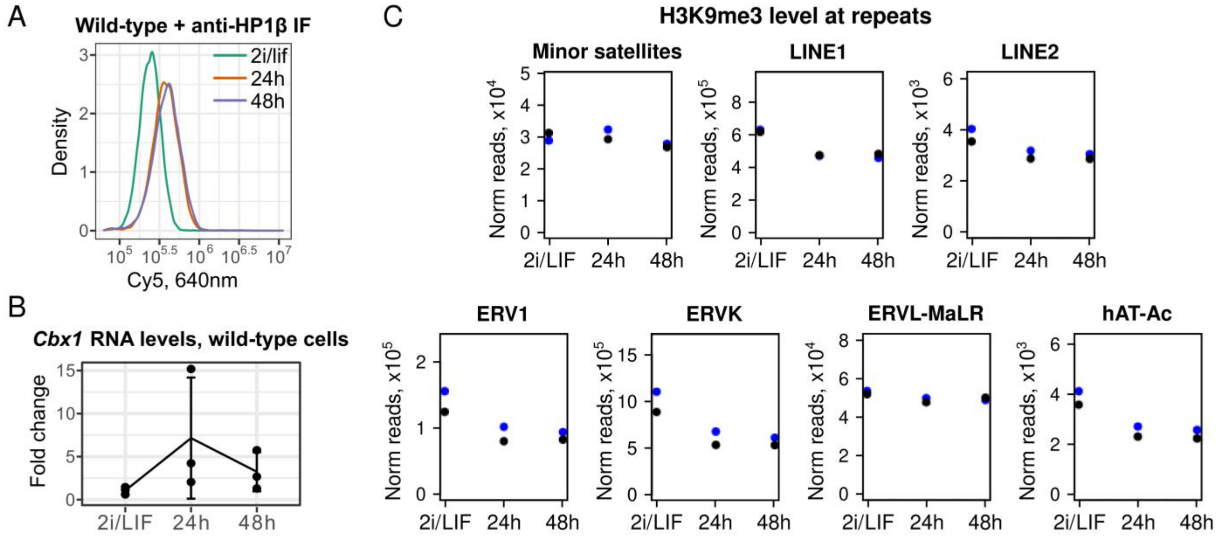


**Figure S6. Characterisation of HP1β and H3K9me3 during exit of mESCs from naïve pluripotency, related to Figure 7.** A. Histograms of the level of fluorescence intensity in single cells in different conditions, measured using flow cytometry after immunolabelling HP1β in wild-type cells. B. *Cbx1* mRNA levels in mESCs and differentiating cells, measured using RT-qPCR (three biological replicates, the error bars show standard deviation). B. Levels of histone H3K9me3 on different types of repetitive sequence, measured using quantitative ChIP-seq (2 replicates).

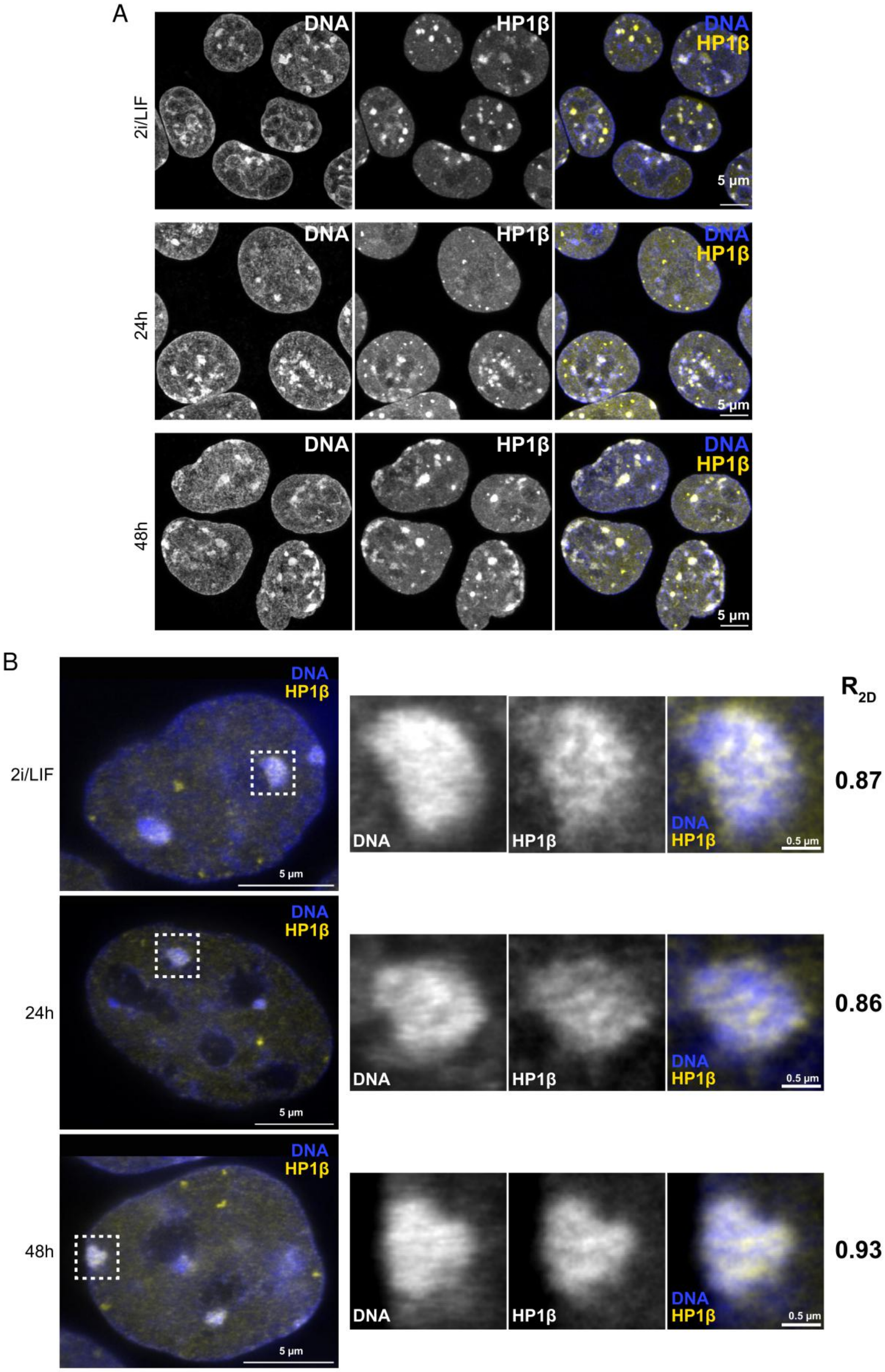


**Figure S7. Detailed Airyscan and STED microscopy images of HP1β and DNA during exit of mESCs from naïve pluripotency, related to Figure 7.** A. Airyscan confocal images of HP1β and DNA in mESCs and during the early differentiation timecourse (maximum projections of z-stacks; the displayed intensities are adjusted for better visibility, and are thus not comparable between images.) Selected cells are also shown in Figure 7C. B. Left, 3D STED images of cells, with areas shown in close-up views in Figure 7F indicated (dashed boxes). Right, the indicated close-up views with the DNA and HP1β channels shown separately, and the corresponding 2D Pearson’s correlation coefficients between the DNA and HP1β distribution within the foci.
