## Supplementary document S2 - Supplementary Note for "HP1*β* and densely packed chromatin form separate microdomains in mouse ES cells, which are reconfigured upon exit from naïve pluripotency"

Let us assume that the HP1β binds to chromatin exclusively through an interaction with the H3K9me3 modification, and that a simple bimolecular reaction takes place: HP1β + H3K9me3 $\rightleftharpoons$ HP1β–H3K9me3. Then the equation for $K_d$ is:

$$\begin{gathered} K_d = \frac{[\mathrm{HP1\beta}_{free}][\mathrm{H3K9me3}_{free}]}{[\mathrm{HP1\beta\text{–}H3K9me3}]} = \\ \frac{([\mathrm{HP1\beta}_{tot}] - [\mathrm{HP1\beta\text{–}H3K9me3}])([\mathrm{H3K9me3}_{tot}] - [\mathrm{HP1\beta\text{–}H3K9me3}])}{[\mathrm{HP1\beta\text{–}H3K9me3}]} = \\ (\frac{[\mathrm{HP1\beta}_{tot}]}{[\mathrm{HP1\beta\text{–}H3K9me3}]} - 1)([\mathrm{H3K9me3}_{tot}] - [\mathrm{HP1\beta\text{–}H3K9me3}]) \end{gathered} \tag{1}$$

Further, if the mode of interaction does not change between different environments, the $K_d$ should also stay constant:

$$\begin{gathered} K_d(\mathrm{eu}) = K_d(\text{DNA-poor, HP1β-rich foci}) = \\ K_d(\text{DNA-rich, HP1β-poor foci}) = K_d(\text{DNA-rich, HP1β-rich foci}) \end{gathered} \tag{2}$$

Naturally, we do not have the necessary measurements to solve these equations in absolute units. However, from SPT experiments, we know the relationship between [$\mathrm{HP1\beta}_{tot}$] and [HP1β–H3K9me3]. Furthermore, from the confocal imaging, we can estimate the relative concentrations of [$\mathrm{HP1\beta}_{tot}$] or [$\mathrm{H3K9me3}_{tot}$] in different environments. What we do not know is the ratio *between* [$\mathrm{HP1\beta}_{tot}$] and [$\mathrm{H3K9me3}_{tot}$]. The estimates for these ratios from our data are listed in Table 1.

**Table 1:** Quantitative relationships between parameters.

| Env | $\frac{[\mathrm{HP1\beta\text{–}H3K9me3}](\mathrm{env})}{[\mathrm{HP1\beta}_{tot}](\mathrm{env})}$ | $\frac{[\mathrm{HP1\beta}_{tot}](\mathrm{env})}{[\mathrm{HP1\beta}_{tot}](\mathrm{eu})}$ | $\frac{[\mathrm{H3K9me3}_{tot}](\mathrm{env})}{[\mathrm{H3K9me3}_{tot}](\mathrm{eu})}$ |
|---|---|---|---|
| **Euchromatin** | 0.45 | 1 | 1 |
| **DNA-poor, HP1β-rich foci** | 0.73 | 2.2 | 2.2 |
| **DNA-rich, HP1β-poor foci** | 0.65 | 1.2 | 1.9 |
| **DNA-rich, HP1β-rich foci** | 0.74 | 2.3 | 3.8 |

By substituting equation (1) into equation (2) for euchromatin a particular environment *env*, and dividing both sides by the concentration of HP1β in euchromatin [$\mathrm{HP1\beta}_{tot}$](eu), we obtain:

$$(\frac{[\text{HP1}\beta_{tot}](\text{eu})}{[\text{HP1}\beta\text{–H3K9me3}](\text{eu})}-1)(\frac{[\text{H3K9me3}_{tot}](\text{eu})}{[\text{HP1}\beta_{tot}](\text{eu})}-\frac{[\text{HP1}\beta\text{–H3K9me3}](\text{eu})}{[\text{HP1}\beta_{tot}](\text{eu})})=$$
$$(\frac{[\text{HP1}\beta_{tot}](\text{env})}{[\text{HP1}\beta\text{–H3K9me3}](\text{env})}-1)(\frac{[\text{H3K9me3}_{tot}](\text{eu})}{[\text{HP1}\beta_{tot}](\text{eu})}\frac{[\text{H3K9me3}_{tot}](\text{env})}{[\text{H3K9me3}_{tot}](\text{eu})}- \tag{3}$$
$$\frac{[\text{HP1}\beta_{tot}](\text{env})}{[\text{HP1}\beta_{tot}](\text{eu})}\frac{[\text{HP1}\beta\text{–H3K9me3}](\text{env})}{[\text{HP1}\beta_{tot}](\text{env})})$$

In equation (3), all quantities but the ratio between the concentration of HP1β and H3K9me3 in euchromatin $\frac{[\text{H3K9me3}_{tot}](\text{eu})}{[\text{HP1}\beta_{tot}](\text{eu})}$ are known. Thus, denoting $\frac{[\text{H3K9me3}_{tot}](\text{eu})}{[\text{HP1}\beta_{tot}](\text{eu})}$ as x, we can write out the following equations for each of the three types of foci.

- $K_d$(eu) = $K_d$(DNA-poor, HP1β-rich foci) gives $(\frac{1}{0.45}-1)(x-0.45)=(\frac{1}{0.73}-1)(x*2.2-2.2*0.73)$, and $x=-0.1$.
- $K_d$(eu) = $K_d$(DNA-rich, HP1β-poor foci) gives $(\frac{1}{0.45}-1)(x-0.45)=(\frac{1}{0.65}-1)(x*1.9-1.2*0.65)$, and $x=0.68$.
- $K_d$(eu) = $K_d$(DNA-rich, HP1β-rich foci) gives $(\frac{1}{0.45}-1)(x-0.45)=(\frac{1}{0.74}-1)(x*3.8-2.3*0.74)$, and $x=0.45$.

While the results from calculations for DNA-rich, HP1β-poor foci and DNA-rich, HP1β-rich foci do not agree precisely, they are not too far off, and the discrepancy can be explained by the limitations of experimental measurements. However, the DNA-poor, HP1β-rich foci stand out in that the corresponding equation has no solution, i.e. the data for this environment is incompatible with the simple bimolecular interaction model. This conclusion makes intuitive sense even from just looking at Table 1: the enrichment in H3K9me3 in DNA-rich, HP1β-poor foci is almost as high as in DNA-poor, HP1β-rich puncta, and yet there is only weak increase in the total HP1β concentration. On the other hand, a similar total concentration and chromatin-bound fraction of HP1β in DNA-rich, HP1β-rich environment is correlated with a much higher density of H3K9me3 sites.

Thus, we conclude that the high proportion of chromatin-bound HP1β and enrichment in HP1β in general in DNA-poor, HP1β-rich foci cannot be solely explained by a higher density of available H3K9me3 within the scope of a simple bimolecular reaction mechanism.
